# Multi-omics and 3D-imaging reveal bone heterogeneity and unique calvaria cells in neuroinflammation

**DOI:** 10.1101/2021.12.24.473988

**Authors:** Zeynep Ilgin Kolabas, Louis B. Kuemmerle, Robert Perneczky, Benjamin Förstera, Maren Büttner, Ozum Sehnaz Caliskan, Mayar Ali, Zhouyi Rong, Hongcheng Mai, Selina Hummel, Laura M. Bartos, Gloria Biechele, Artem Zatcepin, Natalie L. Albert, Marcus Unterrainer, Johannes Gnörich, Shan Zhao, Igor Khalin, Boris-Stephan Rauchmann, Muge Molbay, Michael Sterr, Ines Kunze, Karen Stanic, Simon Besson-Girard, Anna Kopczak, Sabrina Katzdobler, Carla Palleis, Ozgun Gokce, Heiko Lickert, Hanno Steinke, Ingo Bechmann, Katharina Buerger, Johannes Levin, Christian Haass, Martin Dichgans, Joachim Havla, Tania Kümpfel, Martin Kerschensteiner, Mikael Simons, Nikolaus Plesnila, Natalie Krahmer, Harsharan Singh Bhatia, Suheda Erener, Farida Hellal, Matthias Brendel, Fabian J. Theis, Ali Erturk

**Author notes:** These authors contributed equally (Zeynep Ilgin Kolabas, Louis B. Kuemmerle, Robert Perneczky, Benjamin Förstera).

## Abstract

The meninges of the brain are an important component of neuroinflammatory response. Diverse immune cells move from the calvaria marrow into the dura mater via recently discovered skull-meninges connections (SMCs). However, how the calvaria bone marrow is different from the other bones and whether and how it contributes to human diseases remain unknown. Using multi-omics approaches and whole mouse transparency we reveal that bone marrow cells are highly heterogeneous across the mouse body. The calvaria harbors the most distinct molecular signature with hundreds of differentially expressed genes and proteins. Acute brain injury induces skull-specific alterations including increased calvaria cell numbers. Moreover, TSPO-positron-emission-tomography imaging of stroke, multiple sclerosis and neurodegenerative disease patients demonstrate disease-associated uptake patterns in the human skull, mirroring the underlying brain inflammation. Our study indicates that the calvaria is more than a physical barrier, and its immune cells may present new ways to control brain pathologies.

**Graphical Abstract:** 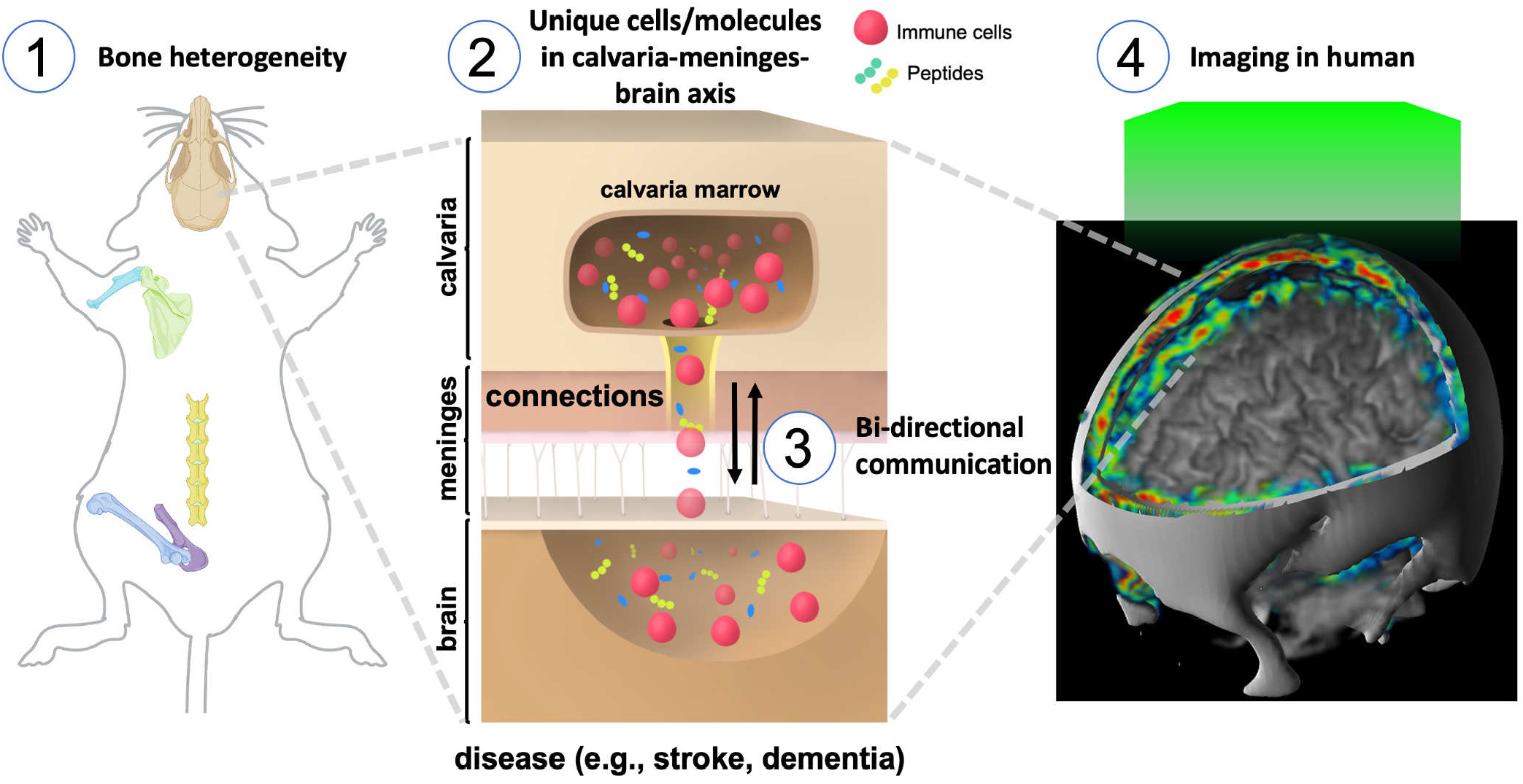

**Highlights:**

1. Bone marrow across the mouse body display heterogeneity in their molecular profile
2. Calvaria cells have a distinct profile that is relevant to brain pathologies
3. Brain native proteins are identified in calvaria in pathological states
4. TSPO-PET imaging of the human skull can be a proxy of neuroinflammation in the brain

**Supplementary Videos can be seen at: http://discotechnologies.org/Calvaria/**

## INTRODUCTION

Neuroinflammation determines major pathological outcomes for many brain diseases, thereby presenting both diagnostic and therapeutic opportunities *(1)*. Many studies not only report increased brain inflammation mediated by microglia activation, but also by diverse somatic immune cells infiltrating the brain from the periphery *(2)*. The meninges surrounding the central nervous system (CNS) serve as a gateway in which peripheral immune cells accumulate and subsequently invade the brain, especially during pathological events *(3)*. Demonstration of skull-meninges connections (SMCs) between the calvaria marrow and meningeal surface of the brain *(4, 5)* suggests that meningeal immune cells might also originate from the calvaria marrow niches. Subsequent studies support the concept of migration of calvaria-originated monocytes, macrophages, and B-cells into dura mater via SMCs *(6–9)*. Yet, the following critical questions remain to be answered *(10)* to understand if calvaria bone can be utilized to monitor and/or control brain inflammation: Are calvaria cells different from other bones in physiological and pathological states? Can brain pathologies induce unique changes in calvaria cells? Finally, what is the relevance for human disease: can we, for example, monitor brain inflammation via imaging of calvaria cells?

The calvaria is positioned as a flat bone on top of the three-layered meningeal membrane surrounding the immune-privileged brain. Here, we investigated whether the calvaria marrow harbors cell populations that could make an important contribution to the brain’s inflammatory state. We used single-cell transcriptomics and mass spectroscopy-based proteomics to compare diverse bones in the mouse body. Our results demonstrate molecular heterogeneity of immune cell populations among the different bones. Notably, the calvaria presented the most distinctive molecular signatures. Acute brain injury induced an increase in the number of calvaria cells with unique profiles that are similar to meningeal cells. Moreover, 18kDa-translocator protein (TSPO) positron-emission-tomography (PET) imaging of Alzheimer’s disease, multiple sclerosis, stroke and 4-repeat tauopathy patients revealed increased tracer uptake in the skull. Notably, the TSPO signal in the skull followed distinct patterns in the different disease processes possibly reflecting differences in the underlying brain pathologies. These findings provide a valuable resource for bone marrow heterogeneity and also reveal that the human skull responds to brain inflammation, which could be monitored as a proxy to follow brain diseases.

## RESULTS

### Bone marrow heterogeneity throughout the body

We set out to assess whether calvaria marrow contains any unique molecular profile of cells among the different bones. We collected bone marrow from three bone types throughout the mouse body (**Fig. 1A**): three flat bones, two long bones, and one irregular bone from cranial to caudal areas: calvaria, scapula, pelvis (ilium), humerus, femur, and vertebra (between thoracic level T5 to lumbar L3), respectively. Dura mater layer of the meninges (further referred to as meninges) and brain samples were also included in the dataset in order to assess the similarity between proximal bone marrow to these compartments.

**Fig. 1.**
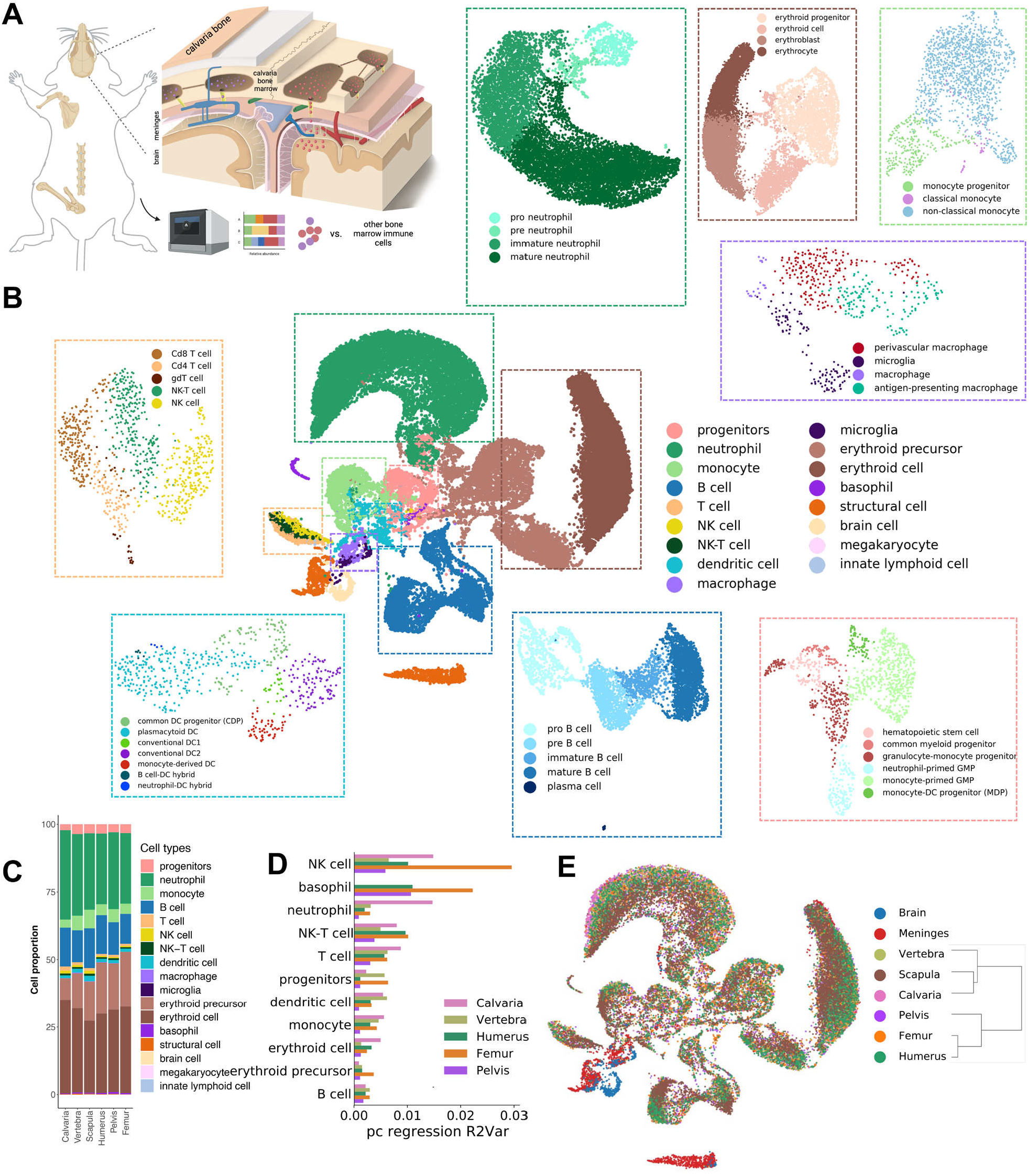
Bones diverge based on transcriptional signature of cell types. (A) Experimental design of single-cell RNA sequencing of calvaria, humerus, scapula, vertebra, pelvis and femur bones, meninges and brain are illustrated with a focus on the complex environment of the calvaria. (B) Cell type umap with fine annotated cell types in the surrounding with matching color. (C) Relative proportions of the coarse cell types are shown in a stacked bar plot. (D) PC regression plot shows how strong each bone’s cell population diverges from the pooled population of other bones by variance explained for each coarse cell type. (E) The region based umap with a dendrogram demonstrates the hierarchical relationship among different bones. (n = 3 animals pooled).

Single-cell transcriptomics analysis of more than 100,000 cells revealed 17 coarse and 54 fine cell types across the bones (**Fig. 1B**). We detected large numbers of neutrophils (∼25%) and erythroid cells (∼30%) in addition to monocytes, B cells, T cells, NK cells, and dendritic cells, among others (**Fig. S1**). Further analysis of the cells using Leiden clustering showed neutrophil-primed or monocyte-primed granulocyte-monocyte progenitors in addition to hybrid cells reported in the literature such as neutrophil-DC cells *(11)* (**Fig. S2A**). Interestingly, we observed high proportions of *Cx3cr1* positive monocytes in all bones, which likely represent the physiological state of the cells *(12, 13)*. Standard cell type proportions were homogeneously distributed among different bones (**Fig. 1C, Fig. S2**). However, principal component regression analysis evidently displayed the transcriptional differences of cell types in different bones (**Fig. 1D, Fig. S2C**). Principal component regression analysis measures how strong one bone’s population diverges from the other bones’ pooled population for each cell type. For example, the calvaria’s neutrophils and monocytes diverged most compared to the neutrophils and monocytes of the other bones. NK cells and basophils of the femur diverged most compared to NK cells and basophils in the other bones. The region-based UMAP displayed a rather homogenous distribution indicating there was no bone-specific abundance for the given cell types (**Fig. 1E**). The dendrogram highlighted the effect of bone type and distance on transcriptomic profiles of the bones. The long bones, femur, and humerus, clustered together with the pelvis. Likewise, the two flat bones situated close to each other, scapula and calvaria, clustered together, and branched with the vertebra (**Fig. 1E**).

### Calvaria cells exhibit distinct migratory signatures

Diverse immune cell populations, including neutrophils, monocytes, and B-cells, can move from calvaria marrow to the dura mater of the meninges via SMCs *(4–7)*. To gain insight into the specific molecular signature of calvaria cells, we analyzed differentially expressed genes (DEGs) and ligand-receptor pairs in all bones (DEGs defined by p < 0.05 and log fold (LF) change > 2 based on two-t-tests, see Methods). We found the highest number of DEGs with around 80 genes, present in the module femur, pelvis, vertebra and humerus (**Fig. 2A**). Most of these genes were expressed in progenitor cells such as pro neutrophils, granulocyte-monocyte progenitors, and erythroid progenitors (**Fig. 2A**). Calvaria displayed the highest number of DEGs (23 genes) among individual bones. The myeloid lineage harbored the highest number of DEGs. Nonclassical monocytes, mature and immature neutrophils, and monocyte-primed granulocyte-monocyte progenitor (GMP)s expressed genes related to the regulation of apoptotic processes and programmed cell death pathways (**Fig. 2B**). Differential expression modules of all bones except calvaria and all bones except vertebra constituted mostly of genes involved in metabolic processes, hinting at a lower metabolic activity in the calvaria and vertebra relative to the other bones. Calvaria-unique DEGs were mostly transcription factors and immediate early genes with a broad range of functions. These genes included *Nr4a1, Nr4a2* (involved in cellular proliferation, apoptosis, and metabolism, T cell regulation) *(14), Egr1* (B-cell development and proliferation) *(15, 16), Osm* (hematopoietic bone marrow and cell mobilization) *(17, 18)*, Ier2 (cell motility and neuronal differentiation) *(19, 20), Apol8* (promoting neuronal differentiation) *(21)* and *Cd14* (Alzheimer’s disease pathogenesis) *(22)*. Lastly, the calvaria exhibited differentially expressed pro- and anti-inflammatory genes, such as *Il1b (23), Ptgs2 (24), Acod (25)*, and *Thbs2 (26)*, some of which are also known to be involved in cell adhesion and migration.

**Fig. 2.**
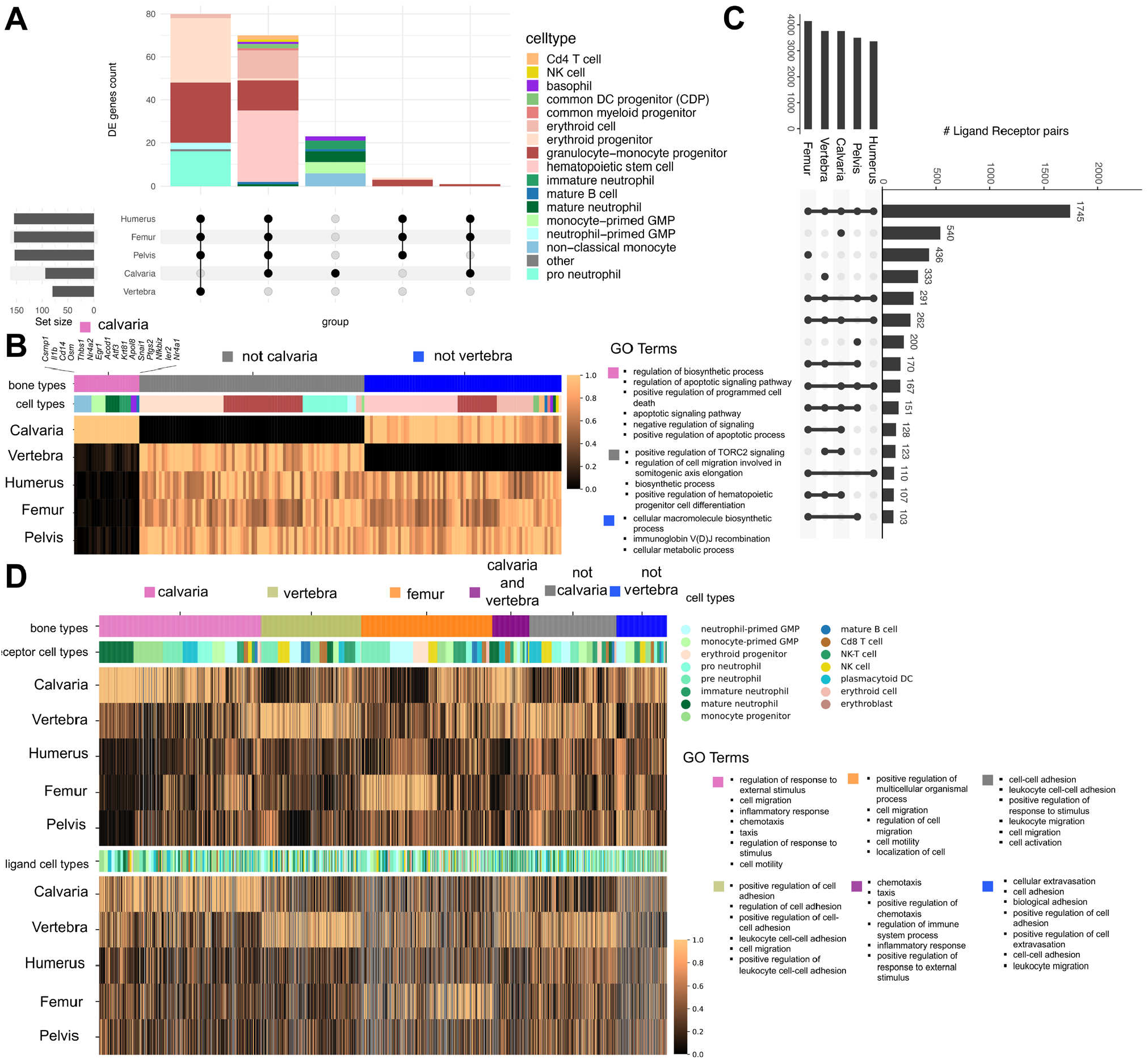
Bone marrow proximal to CNS borders display unique migratory signature. (A) Differentially expressed genes for fine cell types (p<0.05, logFC>2.), display about 80 genes expressed in all bones except for the calvaria in several cell types. Highest number of unique differentially expressed genes are found in the calvaria with >20 genes. *Nra4a1, Nr4a2, Osm, Ptgs2, Atf3* are differentially expressed in more than one cell type. (B) Differentially expressed genes are given in a matrix plot with GO terms associated for each group of genes belonging to only calvaria, all bones except for calvaria and all bones except for vertebra. (C) Ligand-receptor pair analysis among the bones reveal a common interaction module in addition to unique interaction modules in bones, and modules that are present in all bones except for one bone. (D) Ligand-receptor pair analysis showing the ligands (upper) and receptors (lower) in a heatmap. The bar plot with bone names reflect the total number of interaction partners detected for a region, lower bar plot reflect total numbers of detected pairs in a given module. The minimal mean expression among bones is set to zero and the maximal to one. “not *calvaria/vertebra*” means that, that module is present in all other bones except for ‘*calvaria/vertebra*’ in both heatmaps. (n = 3 animals pooled).

We next investigated the ligand-receptor (LR) interactions based on our transcriptomics data using CellPhoneDB and identified several bone-type unique interactions (**Fig. 2C**). We found a core module common to all bones (1745 pairs) mostly involved in cell migration pathways, cytokine production, and regulation of immune system processes such as *Pecam1-Cd1777* and *Cd74-Mif (27)* (**Fig. S2E**). Among unique interactions of individual bones, calvaria (540 pairs) ranked the highest followed by femur (436 pairs) and vertebra (333 pairs) (**Fig. S2E, Fig. S3**). Calvaria showed unique pairs of ligand-receptors mostly in myeloid lineage cells including *Lgals9-Lrp1, Csf1-Slc7a1*, and *Lgals9-Dag1* (involved in migration and adhesion), *Selp-Selpg* (immune cell rolling) *(28)* and *Tnf, Csf1, Il1b*, and *Mif* (pro-inflammatory roles) (**Fig. S2E, Fig. S3**). Vertebra’s unique ligand-receptor pairs revealed terms such as *Icam1* and *Cd44* (adhesion-related), as well as *Cd28* (T-cell related) (**Fig. 2D, Fig. S3**). In conclusion, calvaria displayed the highest number of DEGs and receptor-ligand pairs underlining its distinct molecular profile related to migration and inflammation, especially in the myeloid lineage during homeostatic conditions.

### Calvaria cells respond to pathologies

After identifying bone heterogeneity and unique transcriptome of calvaria cells, we investigated how bone marrows react to a localized brain lesion. To this end, we chose the middle cerebral artery occlusion (MCAo) stroke model in mice *(29)*. In MCAo, the mice first undergo a neck incision to expose the carotid artery before the occlusion of the MCA (**Fig. 3A**). Thus, the sham-operation procedure without MCA occlusion mimics a systemic injury *(29–31)*. We therefore compared MCAo (systemic injury + brain injury), sham (systemic injury alone) and naïve (no operation) to dissect out the specific consequences of brain injury alone.

**Fig. 3.**
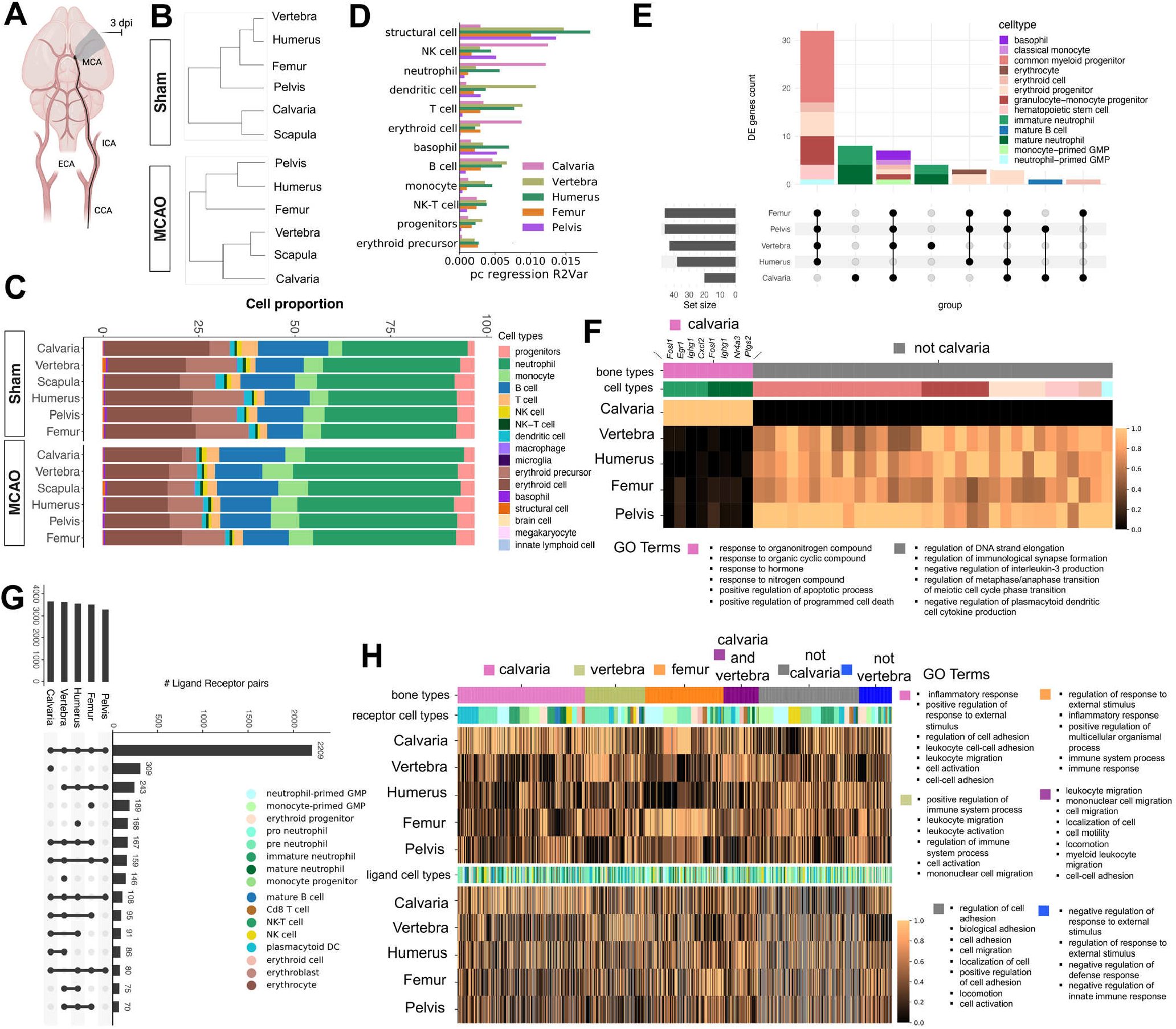
Brain injury instructs shift in transcriptome in which similarity among bones increases. (A) Schematic outline of the MCAo surgery, dpi stands for days post injury. (B) Dendrograms for sham and MCAo conditions. (C) Relative proportions of the coarse cell types after injury: sham and MCAo. (D) Pc regressions reflect how strong each bone’s population diverges from the pooled population of other bones by variance explained for each coarse cell type for both sham and MCAo conditions, combined. (E) Differentially expressed genes in MCAo condition are given. (F) Gene expressions of calvaria unique and calvaria excluding modules are shown in a heatmap with associated GO terms. (G) Ligand-receptor interactions in MCAo condition is demonstrated. There is an overall increase in the common module compared to physiological conditions. (H) The heatmap represents, calvaria, vertebra, femur, calvaria and vertebra common, calvaria excluding and vertebra excluding modules’ ligand (upper) and receptor (lower) expression. Associated biological pathways from GO terms are on the right. The bar plot with bone names reflect the total number of interaction partners detected for a region, lower bar plot reflect total numbers of detected pairs in a given module. The minimal mean expression among bones is set to zero and the maximal to one. (n=3 pooled animals for sham and n=6 pooled animals for MCAo). “not *calvaria/vertebra*” means that, that module is present in all other bones except for ‘*calvaria/vertebra*’ in both heatmaps.

First, we imaged whole mouse bodies at cellular resolution by vDISCO tissue clearing (32) and found that the number of total cells (PI-labeled cells) were increased in the calvaria marrow of mice after stroke compared to controls (**Fig. S4A, Videos S1 and S2**). Next, hierarchical clustering of bone scRNAseq data showed differences between the sham and MCAo, e.g., calvaria clustered on its own in MCAo (**Fig. 3B**). The percentage of neutrophils in both MCAo and sham conditions was increased compared to naïve (**Fig. S1A**). The monocyte proportions were drastically altered, and the B-cell progenitors depleted (**Fig. 3C, Fig. S1B-C**). Pc regression analysis showed that calvaria’s NK cells and neutrophils, and vertebra’s dendritic cells were the most distinct (**Fig. 3D, Fig. S2D**). Differential expression tests revealed significant differences between groups (**Fig. S4D-H**) mostly related to metabolic processes (**Fig. S4F**). Calvaria and vertebra showed common ligand-receptor interaction pairs involved in cell migration which differed from the femur’s cell migration-related pairs (**Fig. S4I,J**).

In the MCAo, while the number of DEGs for all bones decreased, calvaria presented the most DEGs among bones which were expressed in mature and immature neutrophils (**Fig. 3E,F**). Some of these overlapped with the DEGs in naïve conditions, such as *Ptgs2* and *Egr1*, which are involved in inflammatory response *(24, 33)*. Some other unique DEGs included *Nr4a3* (promoting T-cell proliferation and reducing progenitor proliferation) *(14), Cxcl2* (a well-known neutrophil chemoattractant) *(34)* and *Ighg1* (increasing mature B-cells) *(35)*. Ligand-receptor interaction analysis showed that the calvaria had the highest number of distinct pairs (309) also in MCAo, while vertebra had 168, and femur had 189 (**Fig. 3G**). Interestingly, calvaria pairs’ gene ontology results showed terms almost exclusively related to adhesion (**Fig. 3H**). On the other hand, femur for example, had unique pairs more related to inflammatory processes. Calvaria unique interactions involved different subtypes of neutrophils (**Fig. S5**).

### Pro- and anti-inflammatory signatures of calvaria are driven mostly by neutrophils

Next, we investigated the distinct profiles of neutrophils (**Fig. 4**). Examining their developmental trajectories using RNA velocity in its scVelo implementation, we found a subset of mature neutrophils from calvaria cells clustering next to a group of neutrophils found in the meninges (**Fig. 4A**). This observation was also quantified by pseudo-time analysis (**Fig. 4B**). Proportions previously also showed higher mature neutrophils in the calvaria region (**Fig. S1A, sham, and MCAo column**). The representative phase portrait of a calcium-binding gene S100a6 confirmed the validity of our scVelo analysis (**Fig. 4C**). To further investigate the similarity of mature neutrophils in the calvaria and meninges, we performed branching trajectory analysis using partition-based graph abstraction (PAGA) as recently recommended for complex trajectories in a benchmark study *(36)*. We observed a clear distinction between the naïve vs. injury group with the meninges positioned in the middle (**Fig. 4D**). The meningeal neutrophils from the naïve condition connected with almost all bones in the naïve condition, whereas the sham and MCAo meningeal neutrophils indeed connected to the calvaria’s sham and MCAo, revealing a similarity between their late-stage neutrophil population profiles.

**Fig. 4.**
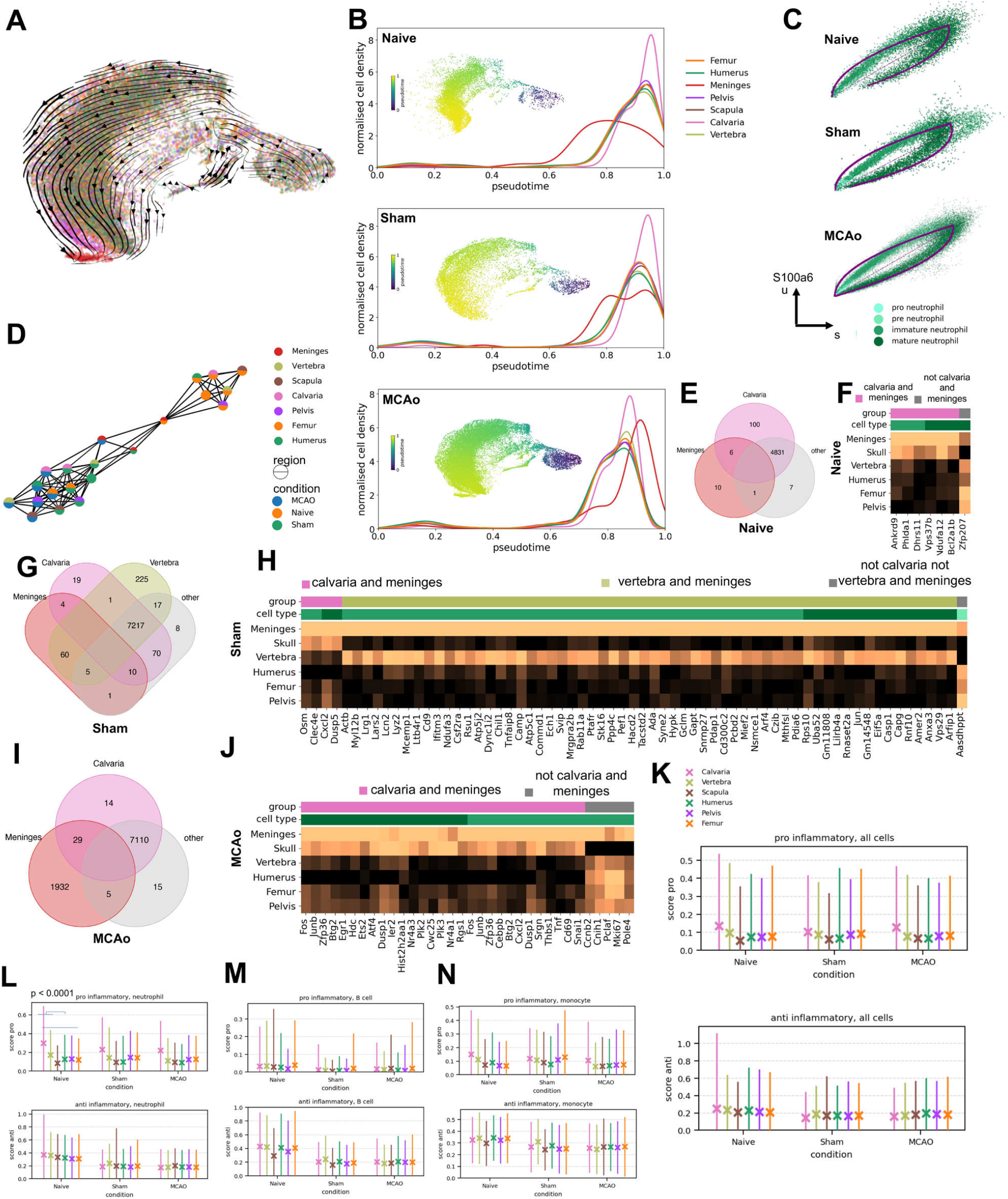
Calvaria neutrophils share transcriptomic signature with meningeal neutrophils after brain injury and pro- and anti-inflammatory scores for bones. (A) Projected developmental trajectory of MCAo neutrophils subset using scVelo. (B) Pseudotime analysis of naïve, sham and MCAo from top to bottom with normalized cell counts in each condition for each region. (C) Phase portrait showing unspliced and spliced counts in neutrophils of gene S100a6 for naïve, sham and MCAo condition respectively. (D) PAGA analysis on the neutrophils subpopulation. (E) Venn diagrams depicting the DE genes among meninges, calvaria and other bones, in naïve condition. (F) Matrix plot shows mean expressions of genes that are up regulated in meninges and in a single other group in naïve condition. (G) Venn diagrams depicting the DE genes among meninges, calvaria and other bones, in sham condition. (H) Matrix plot shows mean expressions of genes that are up regulated in meninges and in a single other group in sham condition. (I) Venn diagrams depicting the DE genes among meninges, calvaria and other bones in MCAo condition. (J) Matrix plot shows mean expressions of genes that are up regulated in meninges and in a single other group in MCAo condition. (K) Mean and standard deviation of pro-(top) and anti-inflammatory (bottom) score over cells of all cell types in naïve, sham and MCAo condition. (L) Mean and standard deviation of pro-(top) and anti-inflammatory (bottom) score over neutrophils in naïve, sham and MCAo condition. (M) Mean and standard deviation of pro-(top) and anti-inflammatory (bottom) score over B cells in naïve, sham and MCAo condition. (N) Mean and standard deviation of pro-(top) and anti-inflammatory (bottom) score over monocytes in naïve, sham and MCAo condition. Significance of the changes in the pro- and anti-inflammatory scores are available in Table S1.

Next, we sought to quantify the pro- vs. anti-inflammatory signatures observed in the calvaria based on the expression of known markers such as *Il6, Il1a, Il1b*, and *Tnf* (pro-inflammatory) and *Il1rn, Tgfb1, Il4, Il10, Il12a*, and *Il13* (anti-inflammatory) *(7, 37)*. Calvaria displayed the highest pro-inflammatory signature among bones in all conditions (**Fig. 4K**). Neutrophils had the highest proinflammatory signature in the calvaria, while B-cells had none (**Fig. 4K-N**). All cells including neutrophils displayed the most anti-inflammatory signature in calvaria in naïve condition (**Fig. 4K-N**). These data imply that the calvaria has both pro- and anti-inflammatory profiles in naïve conditions, which switch to a more pro-inflammatory signature during stroke.

### Protein-level bone heterogeneity corresponds with transcriptomic profiles

Besides the transcriptome, we also investigated the bone heterogeneity using liquid-chromatography mass spectrometry-based proteomics **(Fig. 5A, Fig. S6**). We identified a total of 6159 proteins (**Fig. S6A**). The hierarchical clustering of bone proteome showed similar results to scRNAseq: the calvaria was more distinct than other bones, whereas for example the humerus, femur, and pelvis were closer in PCA space (**Fig. 5B**). Likewise, calvaria proteome branched out separately in hierarchical clustering (**Fig. 5C**). The bone signature in each condition was distinct. The calvaria had 21 proteins uniquely detected in naïve, 20 in sham and 16 in MCAo conditions (**Fig. S6B, Table S3**). Enrichment analysis revealed that cell adhesion, collagen catabolic process, and cell motility terms were frequent in naïve, extracellular matrix related terms in sham conditions. Interestingly, in MCAo, serotonin neurotransmitter release cycle, dopamine neurotransmitter release cycle, presynapse, regulation of neurotransmitter levels, cell adhesion, and biological adhesion terms were detected among others (**Fig. S6B**). No CNS or adhesion related terms were found in unique proteins detected in other bones (data not shown). Additionally, calvaria showed a high abundance of collagen (*Col1a2, Col1a1, Col18a1*), extracellular space (*Nrp2*) and adhesion terms (*Omd, Angptl3*), extracellular space and adhesion terms among bones, which might suggest unique functions besides inflammation (*38*) (**Fig. S6C)**.

**Fig. 5.**
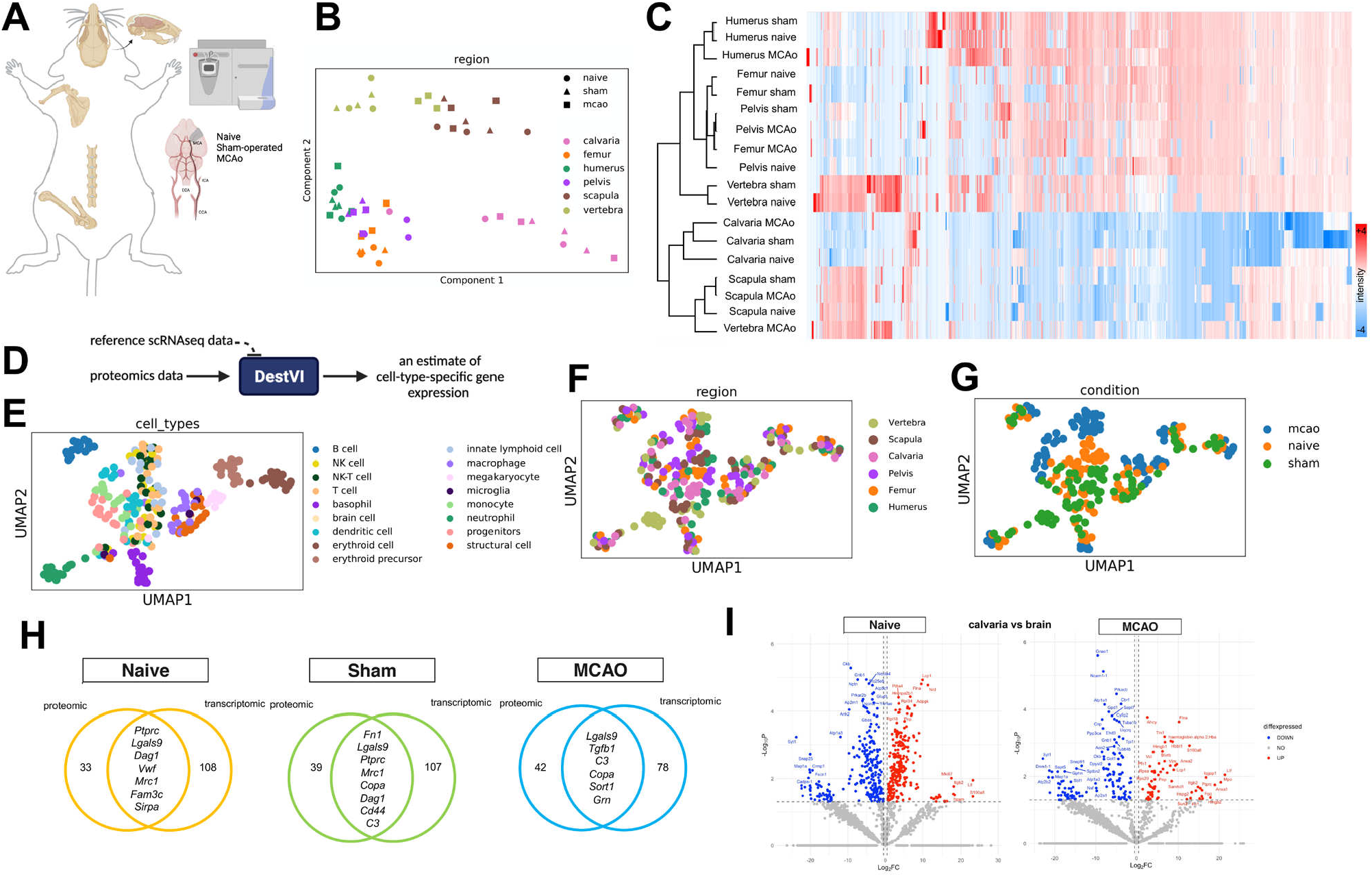
Deconvolution of the cell types in the proteomic data reveals common ligand-receptor pairs highly expressed in the calvaria. (A) Schematic for our proteomics experiment, the experiment was the same as single cell RNA sequencing experiment. (B) Principal component analysis and (C) Hierarchical clustering demonstrate variability among bones over the three conditions: naïve, sham and MCAo. (D) Schematic of DestVI is given, which is applied to our proteomic data to infer an estimate of cell-type specific gene expression. (E) Umap shows coarse cell types after deconvolution with DestVI. (F) Umap shows regions after deconvolution with DestVI. (G) Umap shows conditions after deconvolution with DestVI. (H) Calvaria unique common ligand and/or receptors in scRNAseq data and proteomics data. Volcano plot between calvaria and brain in naïve (left) and MCAo (right) condition (0.05>p, LFC=0.5). (n=3 biological replicates).

Next, we used a deep-learning-based variational inference model, DestVI (*39*), to deconvolute cell types in proteomics data (**Fig. 5D**) under the assumption of correlated protein-expression values per gene. The resulting cell type markers matched with known cell markers such as *Ly6d* and *Igkc* for B-cells, *F10* and *Fn1* for monocytes, *Cd63* and *Ctss* for macrophages, *Mmp9* for neutrophils and *Irf8* for dendritic cells (**Fig. S6E**). The resulting UMAP highly matched with the single-cell transcriptome, which indicates validity of our assumptions for the deconvolution. The neutrophils,

B-cells, and erythroid lineage were separated, and the lymphoid lineage, innate lymphoid cells, NK, NK-T, and T cells were overlayed (**Fig. 5E,F**). We also observed protein groups related to the lymphoid cell and dendritic cell lineage in MCAo but not in other conditions (**Fig. 5G**). Based on this resemblance to transcriptomics analysis, we set out to compare ligand-receptor pairs in calvaria. Seven out of the 40 proteins in naïve, 8 out of the 47 proteins in sham, and 6 out of 48 proteins in MCAo conditions were common to ligand-receptor genes detected in the transcriptomics data (**Fig. 5H**). *Lgals9* was a repeated common protein present in all three conditions and the *Lgals9* and *Dag1* pair was common for the naïve condition.

After MCAo, the correlation between calvaria and brain proteomes slightly increased and showed the highest coefficient of determination (**Fig. S6F**), suggesting a potential exchange of proteins/peptides between calvaria and brain after injury. For example, the number of differentially expressed proteins between calvaria and brain substantially decreased after MCAo (250 vs. 63 upregulated proteins in MCAo calvaria) (**Fig. 5I**). 202 of the upregulated proteins in the skull in naïve condition were no longer upregulated after MCAo. To further examine potential calvaria-brain bi-directional communication, we analyzed the least differentially expressed proteins. We found proteins with increased expression exclusively in the calvaria among bones after MCAo including *Gap43* (involved in presynaptic vesicular function and axonal growth) (*40*) and *Atp1a3* (expressed in neuronal tissue) (*41*) (**Fig. S6I)**. The presence of brain-related proteins in calvaria may suggest a bidirectional calvaria-brain communication.

### TSPO signal in the skull can differentiate inflammatory, ischemic and neurodegenerative CNS diseases

To start investigating the relevance of our findings to human disease, we first used 3D imaging of cleared human post-mortem calvaria samples with dura. Our data showed the presence of SMCs with numerous Iba1+ immune cells (**Fig. 6A**). This suggests potential trafficking of these cells between the calvaria marrow and the dura in humans similar to mice (**Video S3**). We also observed several bone marrow cavities as close as 250 μm to the skull surface, making the cells in the bone marrow cavities more accessible compared to deep brain parenchyma regions (**Fig. 6B**).

**Fig. 6.**
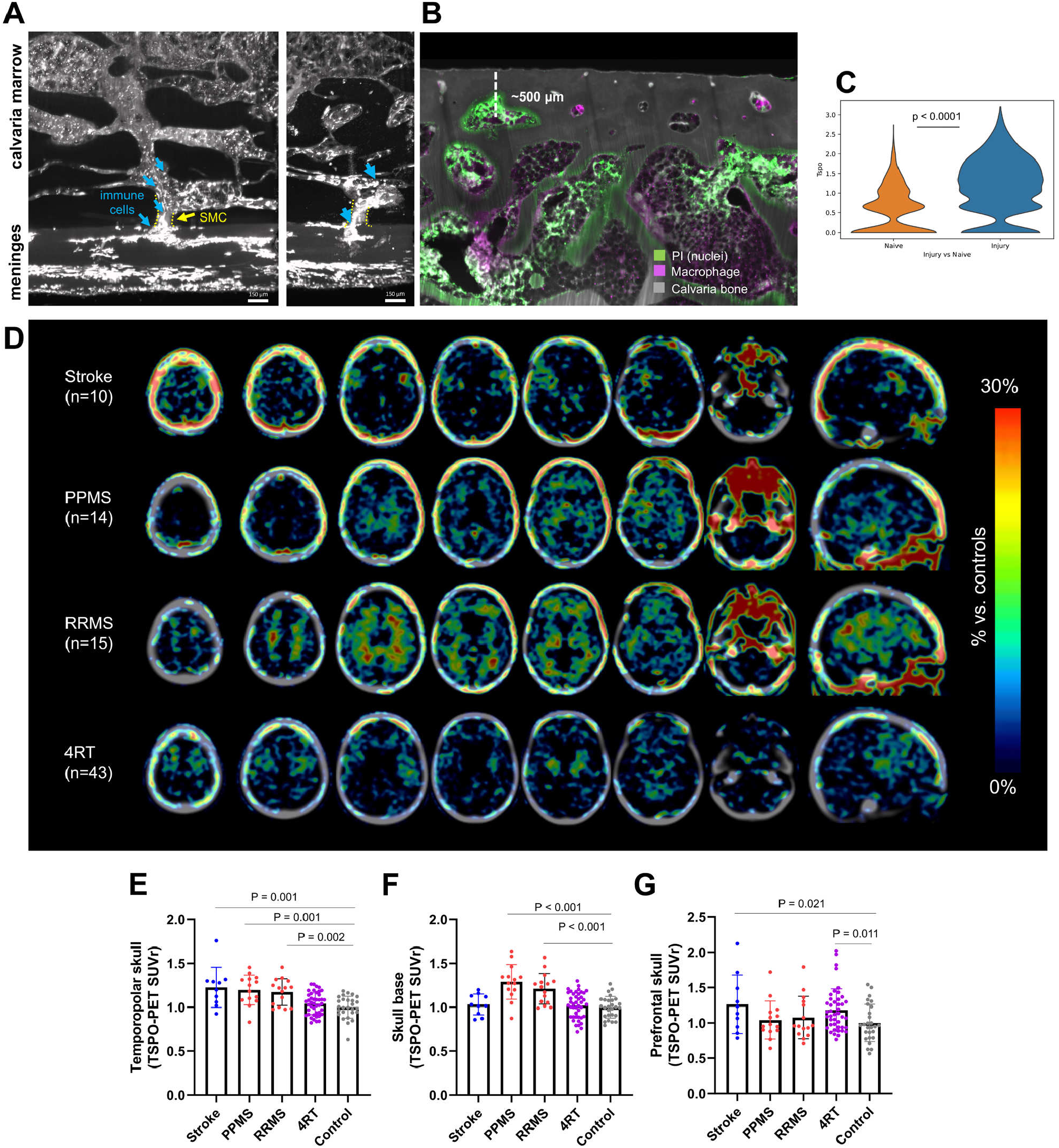
Distinct TSPO uptake patterns are observed in the skull of patients with inflammatory, ischemic and degenerative CNS diseases. (A) Channels connecting calvaria’s bone marrow to the meninges is shown in human calvaria. Iba1+ labeled macrophages are observed in the bone marrow, the channel and the meninges. (B) Human bone marrow labelled for cell nuclei (with PI, in green), macrophage (with Iba1, in magenta) is shown with calvaria bone (autofluorescence). The image depicts the closeness of the bone marrow cavities to the surface of the head to underline the potential for diagnostics and therapeutics of calvaria. (C) Significantly elevated TSPO RNA levels are detected between the naïve vs injury (MCAo + Sham) condition in the skull from the scRNAseq data. (D) Average TSPO-PET signal in stroke, primary progressive multiple sclerosis (PPMS), relapsing remitting multiple sclerosis (RRMS) and 4-repeat tauopathy (4RT) patients. TSPO-PET signal quantifications in skull regions adjacent to different brain regions: (E) temporopolar, (F) skull base, (G) prefrontal area. 1-way ANOVA with Bonferroni post hoc correction. Data were normalized to age-matched controls at the group level prior to analysis. ANOVA included age, sex and TSPO single nucleotide polymorphism as covariates. Significant differences of disease vs. controls are indicated.

After establishing calvaria’s inflammatory response to brain injury, we examined whether this response could be observed in vivo. TSPO is a protein markedly upregulated in the brain during neuroinflammation and is used as biomarker in pre-clinical and clinical research (*42*). We found significantly higher *Tspo* RNA levels in the mouse calvaria in injury conditions compared to naïve (**Fig. 6C**). Next, we assessed whether TSPO imaging of skull would give any hints on underlying brain pathologies in human. To this end, we assessed the TSPO-PET signal in the skull of stroke, relapsing remitting multiple sclerosis (RRMS) (*43*) and primary progressive multiple sclerosis (PPMS) patients as well as 4-repeat tauopathy (4RT) patients (*44*). Our data demonstrated elevated skull inflammation in each cohort of patients with different patterns (**Fig. 6D-G**). In stroke and multiple sclerosis, which show prominent neuroinflammation driven at least partially by immune cell infiltration to the CNS, we observed significantly increased TSPO tracer uptake in the skull covering the temporal pole (**Fig. 6E**). Whereas both RRMS and PPMS patients also showed elevated TSPO uptake in the skull base (**Fig. 6F**) and adjacent bone structures, stroke patients displayed increased TSPO signals in the skull over the prefrontal cortex (*Fig. 6G*). In 4RT patients (movement disorders with motor symptoms and dementia), we observed the increased TSPO signal mainly in the prefrontal region (**Fig. 6G**). These results indicate that TSPO PET imaging of the skull can reveal distinct signal patterns in inflammatory, ischemic and degenerative CNS conditions.

Next, we investigated whether TSPO-PET imaging of skull can also show underlying brain pathology in AD patients, in which neuroinflammation is more subtle compared to stroke and multiple sclerosis. To this end, we assessed TSPO-PET in 52 AD patients compared to 13 healthy controls. We used 3D surface projections on a CT template to show the %-TSPO-PET differences. Our data showed a clear increase in TSPO signal in fronto-parietal calvaria regions of patients with AD, mirroring the increased signal of the underlying fronto-parietal brain regions (**Video S4, white arrow in Fig. 7A,B**). Exemplary section-views on an individual AD patient compared to a healthy control showed a similar TSPO signal increase in the calvaria (**Fig. 7C,D; all subjects quantified in E**). Additionally, the overall TSPO signal was increased in women over men and was negatively associated with age (**Fig. 7F,G**). Our data did not indicate significant associations with clinical severity based on cognitive tests such as Mini-Mental-State Examination (MMSE), the Consortium to Establish a Registry for AD (CERAD) neuropsychological test battery and the Clinical Dementia Rating (CDR) nor any significant associations with specific clinical stages of AD such as in the comparison of the prodromal stage characterized by subjective cognitive decline (SCD) or mild cognitive impairment (MCI) and the AD dementia stage (**Fig. S7A,B**). All AD subgroups displayed a homogenous increase in the calvaria TSPO signal (**Fig. S7A,B**) suggesting that calvaria may prognose all AD subtypes regardless of their stage. We report significant associations between TSPO levels in the calvaria with brain TSPO-PET increases in Braak stage VI regions. Moreover, we report decreased β-amyloid42 but not β-amyloid40 concentration in CSF (with age, sex, MMSE and single nucleotide polymorphism as covariates) (**Fig. S7C-E**). Lastly, we observed a column pattern when assessing regional correlations of calvaria with the brain, which suggests that the brain TSPO, regardless of where it originates, is reflected overall in calvaria TSPO (**Fig. S7F-I**). Overall, these results suggest that the skull can serve as a proxy for monitoring inflammation in the underlying brain regions.

**Fig. 7.**
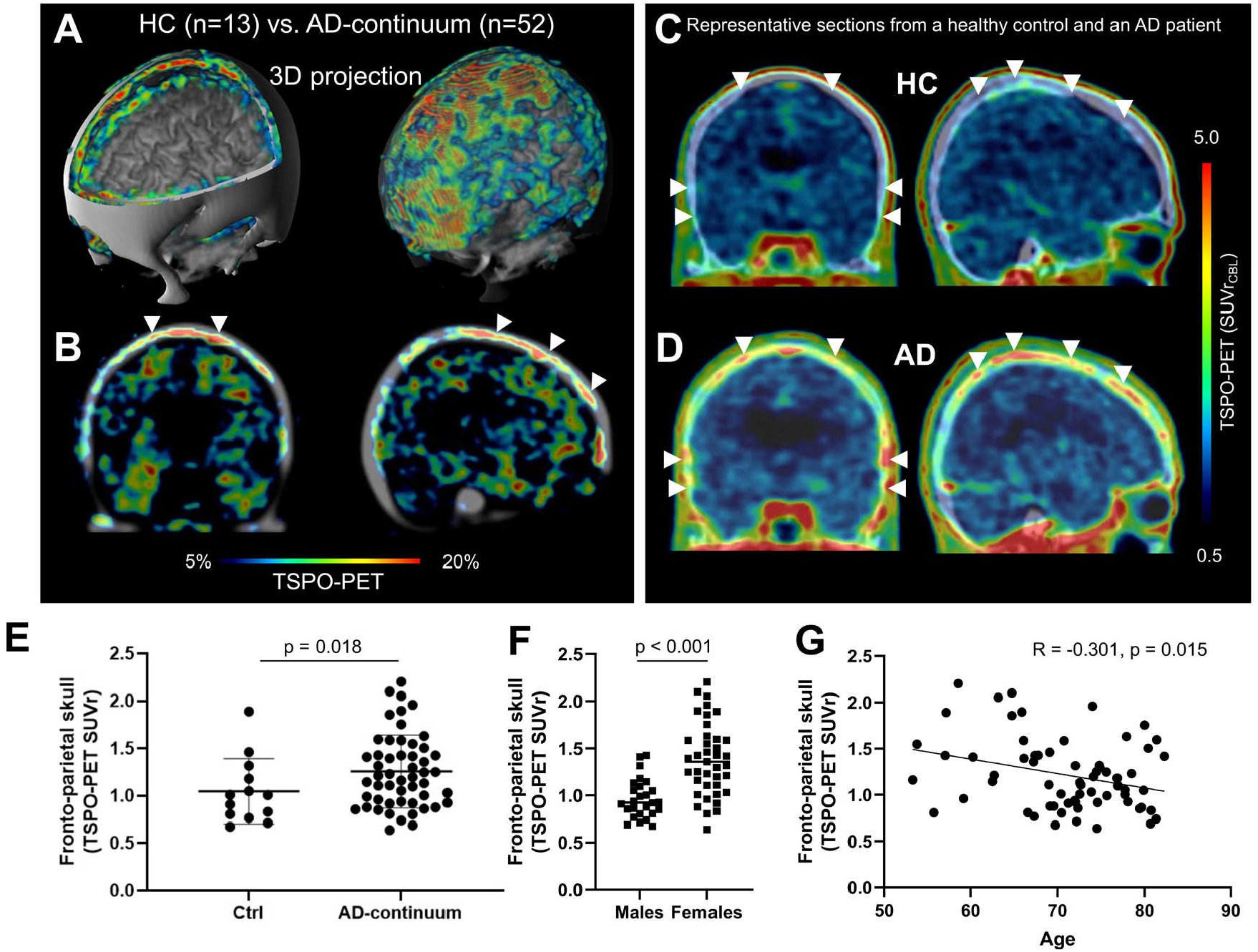
Calvaria inflammation parallels to brain inflammation in Alzheimer patients. (A) 3D surface projection (triple fusion with CT and MRI templates; quadrant cut (left); transparent CT (right) displaying increased activity within skull) shows %-TSPO-PET differences between patients with AD and healthy controls at the group level. Images indicate increased TSPO labelling in the skull in patients with AD when compared to HC pronounced in fronto-parietal skull regions. (B) 2D projection upon CT template shows %-TSPO-PET differences between patients with AD and healthy controls at the group level in skull and brain. Note the strong increases of skull TSPO labelling (indicated by white arrows) in patients with AD when compared to HC which exceed the differences in TSPO labelling of the brain in the same contrast. (C) and (D) Coronal and sagittal slices upon a CT template show TSPO-PET in one healthy control (HC) (57 years old, female, MMSE 29) and in one patient with Alzheimer’s disease (AD; 63 years old, female, MMSE 24). Elevated TSPO labelling in fronto-parietal skull is visualized (white arrows) in the patient with AD when compared to the healthy control. (E) Quantification of fronto-parietal skull signal differences (p=0.018; two-tailed t-test, controlled for age, sex and TSPO-binding single nucleotide polymorphism). (F) Quantification of fronto-parietal skull signal sex differences (p<0.001; controlled for age and TSPO-binding single nucleotide polymorphism) and (G) Quantification of fronto-parietal skull signal age associated patterns (p=0.015, two-tailed t-test, controlled for sex and TSPO-binding single nucleotide polymorphism) among 52 AD continuum patients vs. 13 healthy controls. Data are means ± SD. SUVr = standardized uptake value ratio.

## DISCUSSION

Neuroinflammation has been considered a major pathological event in many CNS disorders. Recently discovered connections between dura and neighboring calvaria marrow suggest that the calvaria could be a new gateway to understand and monitor brain inflammation especially through cells and molecules unique to the calvaria-meninges-brain axis.

### Bone heterogeneity and unique calvaria profile in brain pathologies

The bone marrow contains mesenchymal and hematopoietic stem cells and supplies most cellular components of inflammation, including red blood cells, platelets, and white blood cells. So far, no major differences between different bone marrows have been reported. Our study shows that there is a clear heterogeneity of marrow cells among different bones suggesting the possibility of localized functions in the body through different bones. Bone heterogeneity was more pronounced in the calvaria, which exhibits a different myeloid and B cell gene profile. Genes mostly related to immediate early genes, transcription factors such as the *Nr4a* family (*45*), cell migration and inflammation were highly expressed in the calvaria. We also found various protein/peptide groups related to cell migration, inflammation, and cell stimulation. Moreover, our findings of brain-related proteins in the calvaria marrow suggest that communication along the calvaria-meninges-brain axis might occur in both directions.

Specifically, the neutrophils presented heterogeneous subpopulations reflecting the different stages of maturation (*24, 33, 46*). Analysis of ligand-receptor interaction enabled the identification of interactions essential for migration and adhesion. For example, Lgals9-Dag1 and Lgals9-Lrp1 were detected, which could be specifically targeted to change the course of inflammation in the brain. Yet, currently it is still unclear what fraction of immune cells and under which conditions reach the meninges from the calvaria (*6, 47, 48*) and calvaria independent routes such as blood circulation and choroid plexus (*48*).

### Human calvaria and its involvement in brain pathologies

Here we utilized tissue clearing of large human skull with dura samples (*49*) and demonstrated not only the structure of SMCs at the microscale but also the presence of immune cells inside, suggesting that similar to SMCs in mice, they facilitate immune cell trafficking in humans.

Several studies demonstrated the utility of TSPO-PET imaging of brain to diagnose and monitor neurological diseases such as AD, Huntington disease (HD), amyotrophic lateral sclerosis, Parkinson’s disease, multiple sclerosis and migraine (*50, 51*). Our transcriptomics data also showed increased *Tspo* levels after MCAo-induced stroke in mice. Furthermore, using a third generation TSPO-PET ligand [^18^F]GE-180 (*52*), we observed elevated TSPO-PET signals that showed distinct patterns in the skull of stroke, dementia, multiple sclerosis and movement disorder patients. In AD patients, the TSPO uptake was more pronounced in women and decreased with age maybe due to differences in the skull marrow structure (*53*). Moreover, we observed a stronger global inflammation signal in the human skull compared to the signal from the brain parenchyma, suggesting that skull imaging can be used as an early diagnostic tool for brain pathologies. In AD, the blood-brain-barrier remains mostly intact compared to for example, stroke and multiple sclerosis limiting the penetration of TSPO-PET ligands including [^18^F]GE-180 (*52*) into the brain parenchyma while the skull remains accessible. As calvaria cells are localized very close to the surface, it could be easier and faster to image them by different, for example, optoacoustic imaging (photoacoustic) technologies, which are portable and less costly compared to MRI/PET imaging (*54*). Cloud integration of such portable imaging devices would create early diagnostic and treatment options.

Altogether, our data support the notion that the calvaria uniquely responds to pathologies of the brain. Future research will show if there are similar direct connections from the other ∼200 bones of the human body to the surrounding tissues. Perhaps they can also be utilized to locally monitor and control inflammation and other pathological events such as cancer metastasis.

## METHODS

### Animals

Animal housing and experiments in this work were conducted in agreement to the institutional guidelines (Klinikum der Universität München/Ludwig Maximilian University of Munich, Technische Universitaet Muenchen, Regierung von Oberbayern), after approval of the ethical review board of the government of Upper Bavaria (Regierung von Oberbayern, Munich, Germany), and in accordance with the European directive 2010/63/EU for animal research. The transgenic lines used in this study are C57BJ/J, *LySM*-GFP (Lyz2tm1.1^Graf^, MGI: 2654931) and 5xFAD (B6SJL-Tg(APPSwFlLon, PSEN1*M146L* L286V)6799Vas/Mmjax MGI:3693208) acquired from Charles River and Jackson Laboratory.

### Human skull samples

All donors gave their informed and written consent to explore their cadavers for research and educational purposes, when still alive and well. The signed consents are kept at the Anatomy Institute, University of Leipzig, Germany. Institutional approval was obtained in accordance to the Saxonian Death and Funeral Act of 1994. The signed body donor consents are available on request. The skull sample used in this study comes from a female.

### Middle cerebral artery occlusion (MCAo) model

The MCAo model was used to generate transient cerebral ischemic strokes by introducing an intraluminal filament through the carotid artery of mice anesthetized with isoflurane mixed with 30% O_2_ and 70% N_2_O. To initiate the occlusion the left common carotid artery and interna of the animal were permanently ligated and a silicon capped nylon suture (6/0) was introduced through a cut in the common carotid artery and advanced through the external carotid artery until it reached and obstructed the MCA for 30 minutes. Regional cerebral blood flow was monitored, in the bregma coordinates 2-mm posterior, 5-mm lateral, via transcranial laser Doppler flowmeter from the induction of stroke until 10 minutes after retraction of the filament and reperfusion took place. After the procedure, mice were left for recovery in temperature-controlled cages for two hours in order to minimize the risk of hypothermia. Sham-operated animals were subjected to the same procedure without the insertion of the filament. Body temperatures were kept constant throughout all surgeries with a feedback-controlled heating pad at 37.0 ± 0.5 °C. Animals were then kept in their home cages with facilitated access to water and food whilst being subjected to behavioral tests for three days. Mice were excluded in case of insufficient MCA occlusion (a reduction in blood flow to 15% of the baseline value) or blood flow recovery >80% within 10 min of reperfusion.

### Behavioral experiments - Neuroscore

Neuroscore (*29*) was performed to assess each animal’s general and focal deficits every day. The scoring was composed of general deficits (scores): fur (0 to 2), ears (0 to 2), eyes (0 to 4), posture (0 to 4), spontaneous activity (0 to 4), and epileptic behavior (0 to 12); and focal deficits: body asymmetry (0 to 4), gait (0 to 4), climbing on a surface inclined at 45° (0 to 4), circling behavior (0 to 4), fore-limb asymmetry (0 to 4), compulsory circling behavior (0 to 4), and whisker response to touch (0 to 4). This resulted in a score of 0 to 56 in total; up to 28 from general and up to 28 from focal deficits.

### Perfusion, fixation and tissue preparation

After the mice were anesthetized with a mixture of midazolam, medetomidine and fentanyl (MMF) (1ml/100g of body mass for mice; i.p.), and showed no pedal reflex, they were intracardially perfused with 0.1 M PBS (combined with heparin, 10 U/ml, Ratiopharm). 100-125 mmHg pressure with a Leica 13 Perfusion One system was used for perfusion. PBS ran for 3-4 minutes for single-cell isolation experiment, 5-10 minutes for tissue clearing experiments to let the blood wash out at room temperature. For single-cell isolation experiments, bones were dissected as detailed in the Single cell isolation method section. For the tissue clearing experiments, PBS perfusion was followed by the administration of 4% paraformaldehyde (PFA) in 0.1 M PBS (pH 7.4) (Morphisto, 11762.01000) for 10-20 minutes. After removal of the skin and a washing step with PBS to clean the animal as much as possible, the animals were post-fixed by 4% PFA for the first 24 hours at 4°C and washed three times with 0.1M PBS before processing with the clearing protocol.

### vDISCO whole-body immunostaining, PI labeling and clearing

The detailed protocol of vDISCO was described previously (5). The mouse bodies were placed inside a 300 ml glass chamber (Omnilab, 5163279), to be filled with the appropriate solution regarding the protocol to cover the entire body of the animal (∼250-300ml). A transcardial-circulator system was established in order to allow peristaltic pumping of the solutions (ISMATEC, REGLO Digital MS-4/8 ISM 834; reference tubing, SC0266), with the pressure being set at 180-230 mmHg (50-60 rpm). The tubing was set to allow pumping of the solutions through the heart (attached to a perfusion needle (Leica, 39471024)) into the vasculature with the same entry point used for PBS and PFA perfusion steps described above. The other end of the tube was immersed into the chamber with a loose end to allow suction of the solution into the body. The samples were initially perfused with a decolorization solution (25% of CUBIC reagent 1 (55) which is composed of 25 wt% urea (Carl Roth, 3941.3), 25 wt% N,N,N’,N’-tetrakis (2-hydroxypropyl) ethylenediamine (Sigma, 122262) and 15 wt% Triton X-100 (AppliChem, A4975,1000) in 0.1 M PBS)) for 2 days, refreshing the solutions every 12h. Samples were washed with PBS for 3×2h. Then, decalcification solution (10 wt/vol% EDTA in 0.01 PBS, pH∼8-9, Carl Roth, 1702922685) was perfused for 2 days followed by half a day with permeabilization solution composed of 0.5% Triton X-100, 1.5% goat serum (GIBCO, 16210072), 0.5 mM of Methyl-beta-cyclodextrin (Sigma, 332615), 0.2% trans-1-Acetyl-4-hydroxy-L-proline (Sigma, 441562), 0.05% sodium azide (Sigma, 71290) in 0.01 M PBS. To initiate the PI labeling and boosting, the setup was adjusted. The free end of the perfusion tube was connected to a 0.22 μm syringe filter (Sartorius, 16532) and an infrared lamp (Beuer, IL21) was aimed at the chamber to enable the solution’s temperature to be around 26-28 °C. This setup was then left running for 6 days after the addition of 35 μl of nanobooster (stock concentration 0.5 – 1 mg/ml) and 290 μl of propidium iodide (stock concentration 1 mg/ml) which was added directly into the refreshed permeabilization solution. Next, the body was placed into a 50 ml tube (Falcon, 352070), with the same permeabilization and labeling solution, and an extra 5 μl of nanobooster was added. The tube was then put on a shaker at RT for 2 additional days for labeling. Atto647N conjugated anti GFP nanobooster (Chromotek, gba647n-100) and Propidium iodide (PI, Sigma, P4864), was used to boost the signal from the LysM animals and stain cell nuclei respectively in the study. Then, the animals were placed back into the initial perfusion setup, where washing solution was perfused for 2×12h, which was composed of; 1.5% goat serum, 0.5% Triton X-100, 0.05% of sodium azide in 0.1 M PBS. 0.1 M PBS was used to wash the sample 3×2h. 3DISCO protocol was applied for whole body clearing. The animals were freed from the perfusion system, but kept in glass chambers and placed on top of shakers (IKA, 2D digital) at room temperature inside a fume hood. Glass chambers were sealed with parafilm and covered with aluminum foil along with the 3DISCO application. For dehydration, sequential immersion of tetrahydrofuran (THF) (Sigma, 186562) (50 Vol% THF, 70 Vol% THF, 80 Vol% THF, 100 Vol% THF and again 100 Vol% THF) was applied every 12 hours. Then three hours of dichloromethane (DCM) (Sigma, 270997) immersion for delipidation was followed by indefinite immersion in BABB (benzyl alcohol + benzyl benzoate 1:2, Sigma, 24122 and W213802) solution for refractive index matching.

### Human skull labeling and clearing

Human skull samples were labeled and cleared with modifications to the previously described method (49). The samples (0.5 cm3-1 cm3) were first placed on a shaker and decalcified in 50 ml 20% EDTA solution (pH 8.0) for 15-20 days at 37°C until the bone became soft when touching with fingers. The solution was refreshed every 3-4 days. The samples were then immersed in 30 ml 10% (w/v) CHAPS/ 25% (v/v) NMDEA solution for 3-4 days, refreshing solution once in between, for decolorization and permeabilization followed by 3 times wash in 0.1M PBS. 30 ml serial dilutions (50%-70%-100%-100%) of EtOH were used for dehydration which was followed by DCM/MeOH (2:1 v/v) immersion for one day and ascending serial dilutions of EtOH (70%-50%-0%) to rehydrate the sample, 3-4 h per dilution step. The samples were washed with distilled water and were placed in 30 ml 0.5M acetic acid for 1 day. The samples were washed with distilled water again and placed in 30 ml 4M guanidine hydrochloride for 1 day. Samples were washed with 3 times with 0.1M PBS. Then, each of the samples were incubated with blocking buffer at 37°C for 1 day, followed by incubation with rabbit antibody anti-Iba1 (1:1000) in the antibody incubation buffer for 1 week at 37°C. Next, samples were washed with the washing buffer for 1 day refreshing solution 3 times and then incubated with Alexa 647-conjugated secondary antibodies (1:500) in the antibody incubation buffer for 1 week at 37°C. After washing with 0.1M PBS, propidium iodide (1:100) dye was added to the PBS and incubated for 3 days at 37°C for cell nuclei staining. After labelling, the samples were dehydrated with a series of solutions of EtOH/DiH2O (50%, 70%, 100%, 100% v/v) and delipidated with the DCM solution for 4 hours in each solution, followed by BABB incubation at room temperature for refractive index matching until sample transparency was reached in 1-2 days.

### Light sheet microscopy imaging

Single plane illumination (light sheet) image stacks were acquired using an Ultramicroscope II and Ultramicroscope Blaze (Miltenyi BioTec). The available filter sets were ex 470/40 nm, em 535/50 nm; ex 545/25 nm, em 605/70 nm; ex 560/30 nm, em 609/54 nm; ex 580/25 nm, em 625/30 nm; ex 640/40 nm, em 690/50 nm. The filter sets used to capture LysM signal and the PI labeling were 640/40 nm and 545/25 nm filter sets, respectively. Low magnification whole-body imaging of the LysM mice was performed with Ultramicroscope Blaze, with a1.1x objective, 3×8 tiling with 35% overlap and 6 μm z-step. Exposure time was 120 ms, laser power was 25% and 12% for LysM and PI channels, respectively. The depth of the scans were approximately 13 mm from dorsal and ventral surfaces, which were then reconstructed. The whole head images were taken with an Olympus MVX10 zoom body, which offered zoom-out and -in ranging from 0.63x up to 6.3x. The depth of the scans were approximately 4 mm and the z-step used was 6 μm combined with an exposure time of 200 ms. Human skull samples were imaged with 2×3 tiling, 4x objective 3 μm z-step size and 130 ms exposure time. Laser power was adjusted specifically for each sample in order to prevent signal saturation. Light sheet width was 100% and the thickness was 7 μm to get an N/A of 0.31 μm in all imaging procedures.

### Reconstruction of whole-mouse body and mouse head scans and image quantification

The image stacks were acquired and saved by ImSpector (Miltenyi BioTec) as 16-bit grayscale TIFF images for each channel separately. The whole-body tiled stacks were initially stitched utilizing Fiji/ImageJ to obtain stitching on the xy axis. Next, Vision 4d (Arivis AG) was used to fuse the stacks in the z axis. For heads, one tile stacks were acquired, hence stitching was not necessary. Imaris (Bitplane AG) was used to visualize both whole body and intact mouse heads. For quantification of the mouse heads, manual ROIs were drawn on the frontal and parietal skull bones. The areas above manually selected threshold based on bone marrow coverage were recorded. The quantification graph was analyzed and visualized using Graphpad Prism (version 8.0) (Ordinary one-way ANOVA with multiple comparisons).

### Single cell isolation

Single cell isolation from the calvaria, brain, meninges, humerus, scapula, vertebra, femur and pelvis was done for one animal at a time. Three naïve, six MCAo-operated and three sham operated animals were pooled in threes for single-cell RNA sequencing. Another cohort of three animals for naïve, three animals for sham-operated and three MCAo-operated animals were not pooled and were treated separately for proteomic analysis. These experiments were performed on sham and MCAo animals that had the procedure three days prior to the single cell isolation experiment. Separate equipment was utilized during the isolation to ensure high viability of cells free of contamination. The animals were anesthetized with Isoflurane and then with a Ketamine/Xylazine mixture (0.6 ml Ketamin + 0.3 ml Xylazine + 5.1 ml Saline, 0.2 ml for 20 gr animals). Then animals were transcardially perfused with 10 ml of ice cold 0.1 M PBS. After the blood was rinsed, the calvaria bone, humerus, scapula, vertebra, femur, brain, meninges and the pelvis were dissected and processed by separate people to minimize the time required in order to keep the cell viability to a maximum and conditions comparable for all locations. The isolated cells were processed with 37 °C pre-warmed DMEM (Thermo Fischer, 21013024) with 10% heat inactivated fetal bovine serum (FBS) (Sigma Aldrich, F7524-100ML). For brain cell isolation; the brain was isolated from the calvaria and the rest of the body, then, the cortex was separated and the leptomeninges was removed from the surface, the final sample consisted of the injured region. The sample was placed in 5 ml of trypsin enzyme with 0.05% concentration and incubated in a pre-heated 37 °C water bath for 2 minutes. Following this, the reaction was stopped with 10 ml of 37°C pre-warmed DMEM with 10% heat inactivated FBS, the cells were dissociated by gentle trituration with a 1000μl and 200μl pipette and filtered through 70 μm Falcon™ Cell Strainers (08-771-2). For meningeal cell isolation; after the brain was removed, the meningeal dura layer that was attached to the calvaria bone, was plucked carefully using fine tipped dissection pincers (Dumont #55 Forceps, Dumostar, 11295-51, FST) under a dissection microscope. Leptomeninges was not isolated and therefore is not included in this study. The dissected meninges was placed in 37°C pre-warmed DMEM with 10% heat inactivated FBS solution, shredded with a fine scalpel, gently titrating with a 200μl pipette and filtered through a 70 μm Falcon™ Cell Strainers (08-771-2). For humerus, vertebrae and femur cell isolation; the bone was dissected from the body and the muscles and connective tissue were meticulously cleared off. The bone marrow inside was flushed out to the collection tube with the help of a syringe (Braun, Injekt -F Solo 2-piece Fine Dosage Syringe 1 ml x 100), and further dissection of the bone was performed by fine pincers (Dumont #55 Forceps, Dumostar, 11295-51, FST). The remaining bone was cut in to small pieces and added to the cell mix. This mixture was shortly vortexed with 37 °C warmed DMEM with 10% heat inactivated FBS and filtered with 70 μm (Falcon™ Cell Strainers, 08-771-2). Lastly, for the flat bones, calvaria, scapula and pelvis, after carefully clearing the non-bone parts in the sample i.e., muscles and connective tissue, they were cut in to small pieces (Extra Fine Bonn Scissors, 14084-08, FST), and shortly vortexed and filtered through 70 μm Falcon™ Cell Strainers (08-771-2). After all the samples were ready, they were centrifuged at 4 °C, with 1000 rpm, for 5 minutes. The supernatant of all samples was then discarded and remaining precipitate was put into small 1.5 ml Eppendorf tubes (Eppendorf Safe-Lock Tubes, 1.5 mL, Eppendorf Quality™, 0030120086) after resuspension with DMEM. Cell viabilities and numbers were checked with tryphan blue by an automated cell counter (TC20™ Automated Cell Counter) and controlled by manual counting (Neubauer Cytometry Chamber, MARI0640031).

### scRNA sequencing – 10x Genomics

Samples were used for scRNA-seq if the fraction of dead cells determined by tryphan blue staining was below 50%. Cell suspensions were diluted with PBS, supplemented with 2% FCS, to a final concentration of 1000 cell/ μl and 17.000 cells per sample were loaded onto 10x Chromium Single Cell RNA-seq chips to recover a target cell number of 10.000 cells per sample. scRNA-seq libraries were then generated in three replicates using the 10x Chromium Single Cell 3′ Library & Gel Bead Kit v2 (Chromium™ Single Cell 3’ GEM, Library & Gel Bead Kit v3, 16 rxns) following the manufacturer’s protocol. Libraries were sequenced on an Illumina HiSeq 4000 (150 bp, paired end).

### Single-cell RNA data analysis

#### Count matrix generation

Count matrices were created using CellRanger (v. 3.0.2) aligning reads to the mouse genome mm10 (ensrel97). Spliced and unspliced matrices for RNA-velocity analysis were computed using the velocyto (0.17.17) pipeline.

#### Quality control

Samples were jointly analyzed using scanpy (v. 1.6) and anndata (v. 0.7.5) in Python 3.7. Different quality control filters (56) were used to account for the characteristics of the different samples: In bone samples, all cells with a mitochondrial read fraction higher than 0.2 were removed. In Meninges and Brain samples, thresholds were 0.3 and 0.6, respectively. Further, cells with less than 1000 UMI counts (bone samples) and 500 UMI counts (Meninges, Brain), and more than 50,000 UMI counts were removed. We did not apply a minimum gene filter per cell to retain erythroblasts. All genes expressed in less than 10 cells were removed. To estimate doublets, we used the tool scrublet with a doublet score threshold of 0.1 and removed cells with a higher doublet score. Finally, our filtered dataset contained 128,509 cells expressing 17,040 genes coming from 32 samples.

#### Data preprocessing

To normalize the data with scran (57), size factors were determined as follows: data were first temporarily normalized by total with a target sum of 10,000 per cell followed by log+1-scaling. Then, for each cell, 30 nearest neighbors were computed and data were clustered with leiden clustering at default resolution 1. Small clusters with less than 100 cells were merged with closely related clusters based on the PAGA graph. For PAGA graph calculations we used scanpy’s implementation with default parameters (58). Then, size factors were computed on these clusters and the UMI count data were divided by scran size factors for each cell and log+1-scaled. Then, mitochondrial reads were removed and 4,000 highly variable genes per sample were computed (highly_variable_genes with flavor “cell_ranger” in scanpy). Further, cell cycle scores were computed (score_ genes_cell_cycle in scanpy). To evaluate batch effects, PC regression scores for the variance explained by cell cycle, anatomic region and condition were computed for the full dataset and the MCAo replicates, respectively. PC regression scores were lowest in the condition and replicate covariate, respectively, and therefore no batch effect correction was performed.

### Dendrograms

With scanpy’s dendrogram function scipy’s hierarchical linkage clustering was calculated on a Pearson correlation matrix over regions which was calculated for 50 averaged principal components.

#### Cell type annotations

Cell types were annotated according to a two-step procedure. In a first step a leiden clustering was calculated on the log-normalized data. The leiden clusters were annotated with coarse cell type labels according differentially expressed known markers. In the second step leiden clustering with multiple resolutions were calculated for each coarse cell type. Based on differently expressed known markers, as well as additional information like number of genes (59) and scVelo (60) implementation of RNA velocity (61) the clusters were annotated with fine cell types, and coarse annotations were refined.

#### Variance explained by covariates and PC regression

To quantify how strong cell type populations of each condition/region diverge from the other conditions/ regions the explained variance was calculated by linear regression in PCA space. For conditions pairwise regressions were conducted. For each bone the cell type populations were grouped into the given bone vs the other bones. Scores were only calculated if there were at least 20 cells in each of both groups. 50 principal components were calculated for each cell type. A linear regression on the group variable was calculated for each PC component. R2 scores of the linear regression were multiplied by the eigenvalues of the pc components and normalized by the eigenvalue sum, and finally summed up to the variance explained. We decided to exclude scapula in further downstream analysis because we detected an overall decrease in log counts in this sample (**Fig. S2A**).

#### Combinatorial DE tests

For each gene, two t-tests were calculated to identify if the gene is upregulated in a group of bones. To define the two bone groups for a given gene, bones were ordered by the gene’s mean expression and split in two groups at the highest mean expression gap. The first t-test was conducted on the two groups and the second on the two bones closest to the expression gap. The second test ensures that the expressions of the two closest bones of the two groups are significantly different. The maximal p-value and minimal log fold change of both tests were used to identify DEGs. The chosen thresholds are p < 0.05 and LF change > 2.

#### Ligand receptor (LR) interactions

For each bone ligand receptor interaction, pairs between cell types were calculated with CellPhoneDB’s(*62*) (v 2.1.4) statistical analysis. For a fair comparison between bones, pairs were only calculated on cell types that had at least 10 cells in each bone. An interaction is defined by four variables: ligand, receptor, ligand cell type and receptor cell type.

#### Gene ontology enrichment

Enrichment of Gene ontology (GO) terms for biological processes were calculated using GProfiler (*63*).

#### RNA velocity

RNA velocity (*61*) in its scVelo (*60*)(v 0.2.3) implementation was used as follows: the dataset with spliced and unspliced raw counts was reduced to the given cell type and condition. Then genes were filtered to 2000 genes with at least 20 counts each, and cells were normalized (filter_and_normalize function in scVelo). First and second order moments for velocity estimation with the scVelo’s dynamical model were calculated with default parameters.

#### Pseudotime analysis

Diffusion pseudotime (*64*) was calculated to order cells along the neutrophil maturation trajectory. For naive, sham and MCAo a PCA and neighbors graph were recalculated on the neutrophils population. The default parameters of scanpy’s tl.dpt function was used. As root point we selected the most extreme pro neutrophil cell from the umap. For cell density visualization along pseudotime the cell count was smoothed with a Gaussian kernel according to default parameters of seaborn’s (v 0.11.1) kdeplot function. Densities were normalized for each region separately.

### Sample preparation for proteomics analyses

Sample preparation for total proteome MS analyses were performed as described previously (65). SDC lysis buffer (2% SDC, 100 mM Tris-HCl pH 8.5) was used to lyse the cell pellets at 95°C for 10 min at 1,000 rpm. To shear any remaining nucleic acids, samples were sonicated in high mode (30 sec OFF, 30 sec ON) for 12 cycles (Bioruptor^®^ Plus; Diagenode). Then, protein concentration was determined by BCA and 25 μg of protein was used for further preparation. For reduction and alkylation of the samples, final concentrations of 10 mM TCEP and 40 mM CAA were added. Samples were kept in dark, at 45°C for 10 min with 1,000 rpm shake. Trypsin and LysC were added (1:40, protease:protein ratio) and samples were digested overnight at 37°C, 1,000 rpm shake. Peptides were acidified with 2% TFA, isopropanol with 1:1 volume-to-volume ratio. Custom-made StageTips were used for desalting and purification of the acidified samples were done in, which composed of three layers of styrene divinylbenzene reversed-phase sulfonate (SDB-RPS; 3 M Empore) membranes. Peptides were loaded on the activated (100% ACN, 1% TFA in 30% Methanol, 0.2% TFA, respectively) StageTips, run through the SDB-RPS membranes, and washed by EtOAc including 1% TFA, isopropanol including 1% TFA, and 0.2% TFA, respectively. Peptides were then eluted from the membranes via 60 μL elution buffer (80% ACN, 1.25% NH4OH) and dried by vacuum centrifuge (40 min at 45°C). Finally, peptides were reconstituted in 6 μL of loading buffer (2% ACN, 0.1% TFA) and peptide concentration was estimated via optical measurement at 280 nm (Nanodrop 2000; Thermo Scientific).

### MS sample preparation

Total proteomes were analyzed using an EASY-nLC 1200 (Thermo Fisher Scientific) combined with an Orbitrap Exploris 480 Mass Spectrometer (Thermo Fisher Scientific) and a nano-electrospray ion source (Thermo Fisher Scientific). FAIMS Pro™ with Orbitrap Fusion ™ module was used in analysis of total proteome samples with 2 different compensation voltages (CV) (-50V and -70V).

0.5 μg of Peptides were loaded onto 50 cm HPLC column (75 μm inner diameter, in-house packed with 1.9 μm with C18 Reprosil particles (Dr. Maisch GmbH) at 60°C) and separated by reversed-phase chromatography using a binary buffer system consisting of 0.1% formic acid (buffer A) and 80% ACN in 0.1% formic acid (buffer B).) for MS analysis. 0.5 μg of peptides were loaded on the column at 60°C, and separated with a 120 min gradient (5-30% buffer B over 95min, 30-65% buffer B over 5min, 65-95% over 5 min, and wash with 95% buffer B for 5 min) at a flow rate of 300 nL/min. MS data were acquired using a data-dependent MS/MS cycle time scan method. The time between the master scans were 1 second.

Full MS scans were acquired in 300–1650 m/z range with a normalized AGC target %300 at 60,000 resolution and 25 ms maximum injection time. Precursor ions for MS/MS scans were fragmented by higher-energy C-trap dissociation (HCD) with a normalized collision energy of 30%. MS/MS scans were acquired 15,000 at m/z resolution with a normalized AGC target 100%, and a maximum injection time of 28 ms. Expected LC peak width was 120 s and desired apex window was set to 30%.

### Proteomic data processing

The raw data were processed with MaxQuant version 1.6.14.0. Default settings were used if not stated otherwise. FDR 0.1% was used for filtering at protein, peptide and modification level. As variable modifications, acetylation (protein N-term) and oxidized methionine (M), as fixed modifications, carbamidomethyl (C) were selected. Trypsin/P and LysC proteolytic cleavages were added. Missed cleavages allowed for protein analysis was 3. “Match between runs” and label free quantitation (LFQ) were enabled and all searches were performed on the mouse Uniprot FASTA database (2019). Perseus (version 1.6.14.0) was used for bioinformatics analysis. Gene Ontology (GO), UniProtKB and the Kyoto Encyclopedia of Genes and Genomes (KEGG) annotations were included in the analysis. Number of proteins per sample graph visualized using Graphpad Prism (version 8.0). Quantified proteins were filtered for at least two valid values among three biological replicates in at least one group (each group represents each region in each condition) for total proteome analyses. Missing values were replaced using minimum value for each sample’s expression level. Proteins that are expressed significantly different from each other were determined by ANOVA. Permutation based FDR (0.05) and S0-parameter (0.1) were used for truncation. Hierarchical clustering of significant proteins was done by applying Euclidean as a distance measure for row clustering after normalization of median protein abundances of biological replicates by z-score. The correlation plots were computed using Pearson correlation with average linkage.

### Integration of single-cell data and proteomics data

To infer abundance of cell type-specific expression profiles in the samples, we utilized a newly developed variational inference model from scvi-tools (https://scvi-tools.org) namely DestVI (Deconvolution of Spatial Transcriptomics profiles using Variational Inference) (39), where the single cell transcriptomics data was leveraged to deconvolve the bulk spots of the spatial transcriptomics data. We adapted this model for our bulk proteomics data, aiming to assign protein expression to particular cell types after deconvolution. The underlying noise model assumption was that proteomics gene-expressions follow negative binomial distribution as the spatial transcriptomics data (*66, 67*). This assumption can be further studied and the model can be adapted to a different distribution that better describes the proteomics data.

We assumed that since single-cell transcriptomics data and the proteomics data are coming from the same bone although from different biological replicates, are corresponding to one another. As a result, we inferred cell type specific protein profiles from bulk proteomics data and applied the same downstream analysis as for the transcriptomic data (See Methods Variance explained by covariates, Combinatorial DE tests and Ligand receptor interactions for further detail.)

As single-cell data was used as a prior for the deconvolution, we are aware that it could introduce a potential bias. For example, the separated vertebra cluster on region UMAP is only from the sham condition, thus can be considered an artifact rather than a biological signal (**Fig. 5F**).

### Human TSPO-PET imaging

Ten patients with stroke, 29 patients with multiple sclerosis (15 with relapsing remitting multiple sclerosis and 14 with primary progressive multiple sclerosis), 43 patients with 4R tauopathies, 52 patients with AD and 27 age- and sex-matched individuals without objective memory impairment and with intact motor function were available for calvaria analysis of TSPO-PET. In one set of analyses stroke, MS and 4R tauopathy patients were compared with controls, while the AD cohort, for which additional biomarkers were available, was analysed separately. All participants were scanned at the Department of Nuclear Medicine, LMU Munich, using a Biograph 64 PET/CT scanner (Siemens, Erlangen, Germany). Before each PET acquisition, a low-dose CT scan was performed for attenuation correction. Emission data of TSPO-PET were acquired from 60 to 80 minutes (*44, 68*) after the injection of 187 ± 11 MBq [18F]GE-180 as an intravenous bolus, with some patients receiving dynamic PET imaging over 90 minutes. The specific activity was >1500 GBq/μmol at the end of radiosynthesis, and the injected mass was 0.13 ± 0.05 nmol. All participants provided written informed consent before the PET scans. Images were consistently reconstructed using a 3-dimensional ordered subsets expectation maximization algorithm (16 iterations, 4 subsets, 4 mm gaussian filter) with a matrix size of 336 × 336 × 109, and a voxel size of 1.018 × 1.018 × 2.027 mm. Standard corrections for attenuation, scatter, decay, and random counts were applied.

To compare different patient cohorts we used harmonized data from different PET imaging studies: All patients with acute ischemic stroke (n=10) were recruited from the ongoing ICARUS study which included a TSPO-PET up to 10 days after stroke onset. Inclusion criteria were an age >50 years, patients with acute ischemic stroke (time frame: <72 hours) as defined by an acute focal neurological deficit in combination with a corresponding infarct as documented by a diffusion weighted imaging (DWI)-positive lesion on brain MRI. Presence of an infarct involving the cortex or a strictly subcortical infarct; fluent in German language; in patients with cognitive impairment: availability of an informant; written informed consent; willingness to participate in study assessments including follow-up. Exclusion criteria were prior history of stroke, multiple infarcts, infratentorial infarcts affecting the brain stem or cerebellum; immunomodulatory therapies within the last 3 months; chronic inflammatory disease; infectious diseases (< 7 days prior to stroke). PET acquisition and PET data analyses of the stroke cohort (ethics-application: 19-0428) were approved by the local institutional ethics committee (LMU Munich) and the German radiation protection (BfS-application: Z 5 - 22464/2019-163-G) authorities. All patients with multiple sclerosis were investigated during observational studies. We included all baseline scans of therapy naïve patients with primary progressive multiple sclerosis (n=14) and patients with relapsing remitting multiple sclerosis (n=15; previously published in (*43*)) regardless of therapy regimes. However, patients who received steroid therapy < 4 weeks prior to PET as well as patients with additional CNS pathologies were excluded a priori. PET acquisition and PET data analyses of the multiple sclerosis cohort (ethics-application: 601-16) were approved by the local institutional ethics committee (LMU Munich) and the German radiation protection (BfS-application: Z 5 – 22463/2 – 2015 -006) authorities. The 4R-tauopathy cohort (*44*) was composed of patients with possible or probable β-amyloid negative corticobasal syndrome (n=29) and patients with possible or probable progressive supranuclear palsy Richardson syndrome (n=14) according to Armstrong Clinical Research and Movement Disorders Society criteria respectively. Detailed inclusion and exclusion criteria were published elsewhere (*44*). One case was excluded due to cropped skull. PET acquisition and PET data analyses of the 4R-tauopathy cohort (ethics-applications: 17-569 & 17-755) were approved by the local institutional ethics committee (LMU Munich) and the German radiation protection (BfS-application: Z 5 - 22464/2017-047-K-G) authorities. A total of 27 healthy controls deriving from the different cohorts were included to cover the whole age range of patients.

The AD cohort was composed of 9 cases with subjective cognitive decline due to AD, 13 cases with mild cognitive impairment due to AD, 18 cases with AD dementia, and 12 cases with corticobasal syndrome, dementia and underlying AD. Initial results of brain TSPO labeling in this cohort are published elsewhere (*69*). Participants were enrolled in the interdisciplinary AD study “Activity of Cerebral Networks, Amyloid and Microglia in Aging and AD (ActiGliA)”. In the AD cohort and its controls, Aβ-PET was performed in all participants using [18F]flutemetamol (*70*). PET acquisition and PET data analyses of the AD cohort (ethics-applications: 17-569 & 17-755) were approved by the local institutional ethics committee (LMU Munich) and the German radiation protection (BfS-application: Z 5 - 22464/2017-047-K-G) authorities. Emission data of Aβ-PET were acquired from 90 to 110 minutes after injection of 188 ± 10 MBq [18F]flutemetamol. Aβ-PET was assessed by a visual read (one expert reader), and the decision of Aβ-positivity/negativity was supported by a software-driven approach implemented in HERMES Gold (V4.17, HERMES Medical Solutions AB, Stockholm, Sweden). One positive evaluated target region (frontal, temporal, parietal, posterior cingulate) defined the scan as positive.

All TSPO-PET data were analyzed using PMOD. Spatial normalization was performed to a tracer specific templates in the Montreal Neurology Institute (MNI) space which was acquired via MRI-based spatial normalization. All images were normalized by cerebellar grey matter scaling (defined by the Hammers atlas (*71*)) prior to analysis and a standardized-uptake-value (SUV) analysis served for pseudo-reference tissue independent validation.

For stroke, multiple sclerosis and 4R tauopathy patients we defined three target regions based on a voxel-wise exploratory analysis: temporopolar skull (comprising 18 cm^3^), skull base (comprising 97 cm^3^), and prefrontal skull (comprising 7 cm^3^). All regions were semi-automatically delineated using the human CT template available in PMOD. Region-based PET values were normalized to a composition of values of exactly age-matched (≤ 1 year difference) controls at the group level. Voxel-wise differences (% vs. age-matched controls) were calculated to allow a volume-of-interest independent validation of elevated skull tracer binding in all patient groups. Following the region-based approach, we used compositions of exactly age-matched controls for this calculation.

For the AD cohort, TSPO labeling in the calvaria was obtained in each participant from a large fronto-parietal volume-of-interest (comprising 66 cm^3^), which was semi-automatically delineated using the human CT template available in PMOD. Posterior and frontal calvaria was spared to avoid signal spill-over from sinuses and extracranial structures. Furthermore, we used a Brainnetome (*72*) atlas-based classification of cortical brain regions and corresponding calvaria regions to test for regional calvaria-brain associations. To this end, we increased the dimension of the atlas by a factor of 1.2 and we delineated all volumes-of-interest that were represented in the calvaria as defined by the CT template (≥50% of voxels included). This approach resulted in 64 individual calvaria-brain region pairs. TSPO labeling of the calvaria was compared between AD patients with β-amyloid pathophysiology (AD) and β-amyloid negative controls. Voxel-wise differences were calculated to allow a volume-of-interest independent validation of elevated calvaria tracer binding in patients with AD. TSPO labeling of the calvaria was correlated with age, sex, and cognitive testing (MMSE, CERAD, CDR) as well as with β-amyloid levels in CSF. Calvaria-brain associations of TSPO-PET were tested for the global calvaria volume-of interest with Braak stage and β-amyloid related composite brain regions. Furthermore, calvaria-brain associations were tested by a correlation matrix of the predefined 64 volume-of-interest pairs. Single region increases in patients with AD vs. healthy controls were correlated between calvaria and brain regions.

### Statistics for human TSPO-PET imaging

Group comparisons of VOI-based PET results between patient groups with mixed neurological disorders and controls (n=5 groups) were assessed by 1-way ANOVA and Bonferroni post hoc correction for multiple comparisons using IBM SPSS Statistics (version 22.0; SPSS). All data were controlled for age, sex and the TSPO single nucleotide polymorphism at the individual subject level.

Group comparison of Human TSPO-PET results between controls and AD patients were assessed by a two tailed t-test in SPSS Statistics (version 22.0; SPSS), controlled for age, sex and the TSPO single nucleotide polymorphism. For correlation analyses, Pearson coefficients of correlation (R) were calculated. A threshold of P less than 0.05 was considered to be significant for rejection of the null hypothesis.

### Code availability

All code used in this study can be found as jupyter notebooks in the project github repository: https://github.com/erturklab/skull_immune.

### Data availability

Patient source file human TSPO-PET imaging study can be found under Supplementary Information. Single-cell sequencing data raw counts matrices and annotation are available via NCBI’s GEO. The mass spectrometry data have been deposited to the ProteomeXchange Consortium via the PRIDE *(73)* partner repository with the dataset identifier PXD030250.

## ACKNOWLEDGEMENTS

This work was supported by the European Research Council (ERC), Vascular Dementia Research Foundation, Deutsche Forschungsgemeinschaft (DFG, German Research Foundation) under Germany’s Excellence Strategy within the framework of the Munich Cluster for Systems Neurology (EXC 2145 SyNergy, ID 390857198), grant DI-722/16-1 (project ID: 428668490) and a dedicated grant to M.B. (BR4580/1-1). H.M. and Z.R. are supported by the China Scholarship Council (CSC) (No. 201806780034) and M.M is supported by Turkish Ministry of Education. H.M., Z.R. and M.M. are members of Munich Medical Research School (MMRS) at the LMU Munich. A.E and Z. I. K. are members of the Graduate School of Systemic Neurosciences at the Ludwig Maximilian University of Munich. S.P. and C.P. are funded by Lüneburg Heritage Foundation. We thank Doris Kaltenecker for discussions, Mihail I. Todorov for discussion on image analysis and Alex Nazlidis for contribution to Fig 1a. The illustration of the experimental pipeline in Fig. 1, 3 and 4 was created with BioRender. com. The work of Mat.B., B-S.R. and R.P. was supported by the Hirnliga e.V. (Manfred-Strohscheer-Stiftung) and the Alzheimer Forschung Initiative e.V. (project ID 19063p). GE Healthcare made GE-180 cassettes available through an early-access model. The ICARUS study was supported by the DFG immunostroke project through FOR 2879 (project number 428668490). N.K. and O.S.C. are supported by the DFG Emmy Noether Programme (KR5166/1-1). J.L. is supported by the BMBF (FKZ: FKZ161L0214B, ClinspectM).

## AUTHOR CONTRIBUTION

Z.I.K. designed, performed and analyzed most of the experiments. L.B.K., Mar.B., Z.I.K. and F.J.T. performed the single-cell RNA sequencing bioinformatic analysis. R.P. provided ActiGliA clinical data and TSPO-PET scans. B.F. designed, coordinated and performed experiments in the initial phase of the project and performed MCAo and sham surgeries. H.S.B co-organized experiments in the initial phase of the project. O.S.C., Z.I.K. and N.K. performed mass-spectrometry experiment and analysis. M.A. performed the mass-spectrometry analysis and deconvolution of the proteomics data. Z.I.K., B.F., H.S.B., Z.R., H.M., M.M., K.S., F.H., Ig.K., N.P., S.B.G. and O.G. contributed to sample collection for RNA sequencing and mass spectrometry. M.S., In.K., H.L. performed cDNA synthesis and library construction for single-cell RNA sequencing. N.L.A., M.U., J.G., S.K., C.P., A.K., K.B., J.L., C.H., M.S., M.D., J.H., T.K., M.K., R.P., Mat.B. organized, recruited, and conducted the human TSPO-PET study. G.B., L.M.B. S.H., J.G. and Mat.B. analyzed human TSPO-PET scans. A.Z. produced 3D projections. H.S. and I.B. provided human skull sample. S.Z. processed and imaged the human skull sample. F.H., S.E., H.S.B., and A.E. supervised the project. Z.I.K. and S.E. provided the first draft of the manuscript, F.H., H.S.B., L.B.K. helped write the manuscript. A.E and Z.I.K cowrote the final manuscript. A.E conceived the project and led all aspects of the project. All authors critically reviewed, contributed to and approved the final manuscript.

## Video legends

**Video S1**. vDISCO whole-body cleared LysM mouse demonstrates the distribution of monocutes and neutrophils in the whole body after MCAo.

**Video S2**. vDISCO cleared LysM mouse head after MCAo has high cell instensity as well as LysM+ immune cells in parietal and frontal skull.

**Video S3**. Iba+ labeled human skull+meninges (dura) sample displays immune cells in the skull-meninges connections

**Video S4**. TSPO-PET signal from an Alzheimer’s patient shows high TSPO signal coming from the calvaria.

## Tables

**Table S1**. Number of cell counts per cell type and condition

**Table S2**. Significance of changes in pro- and anti-inflammatory scores for bones

**Table S3**. Calvaria only genes captured from proteomics samples for each condition

**Table S4**. Patient source file for human TSPO-PET imaging data

**Fig. S1.**
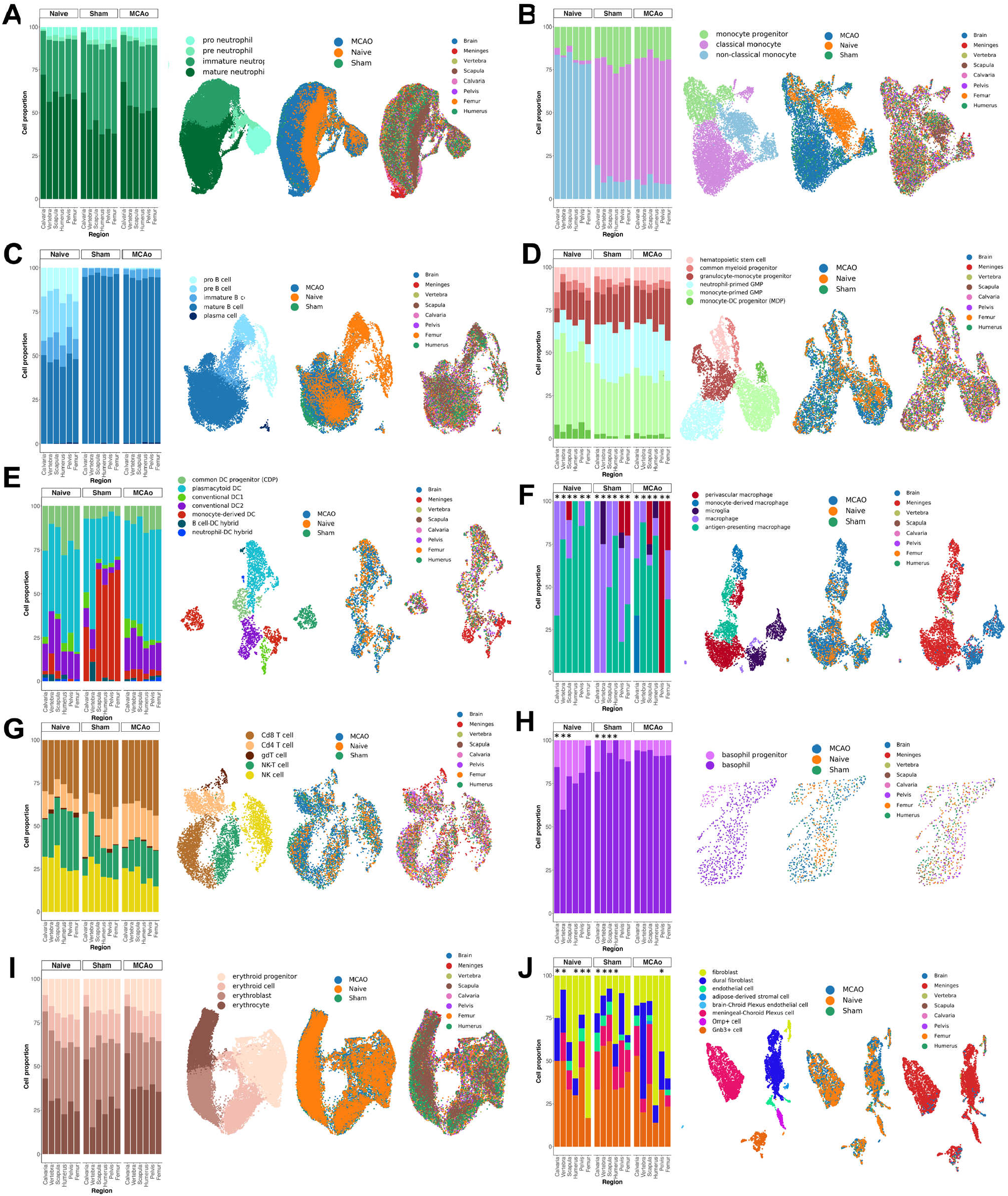
Proportions and umap of fine cell-types over all conditions. Coarse cell types are shown separately with their fine cell type proportion over three conditions, and their umap distribution for the cell type, condition and region. Shown coarse cell types include (A) neutrophils, (B) monocytes, (C) B cells, (D) progenitors, (E) dendritic cells, (F) macrophages, (G) T and NK cells, (H) basophils, (I) erythroid cells (J) structural cells. Regions with cell types less than 20 cells are indicated with a * on their relative positions in the bar plots.

**Fig. S2.**
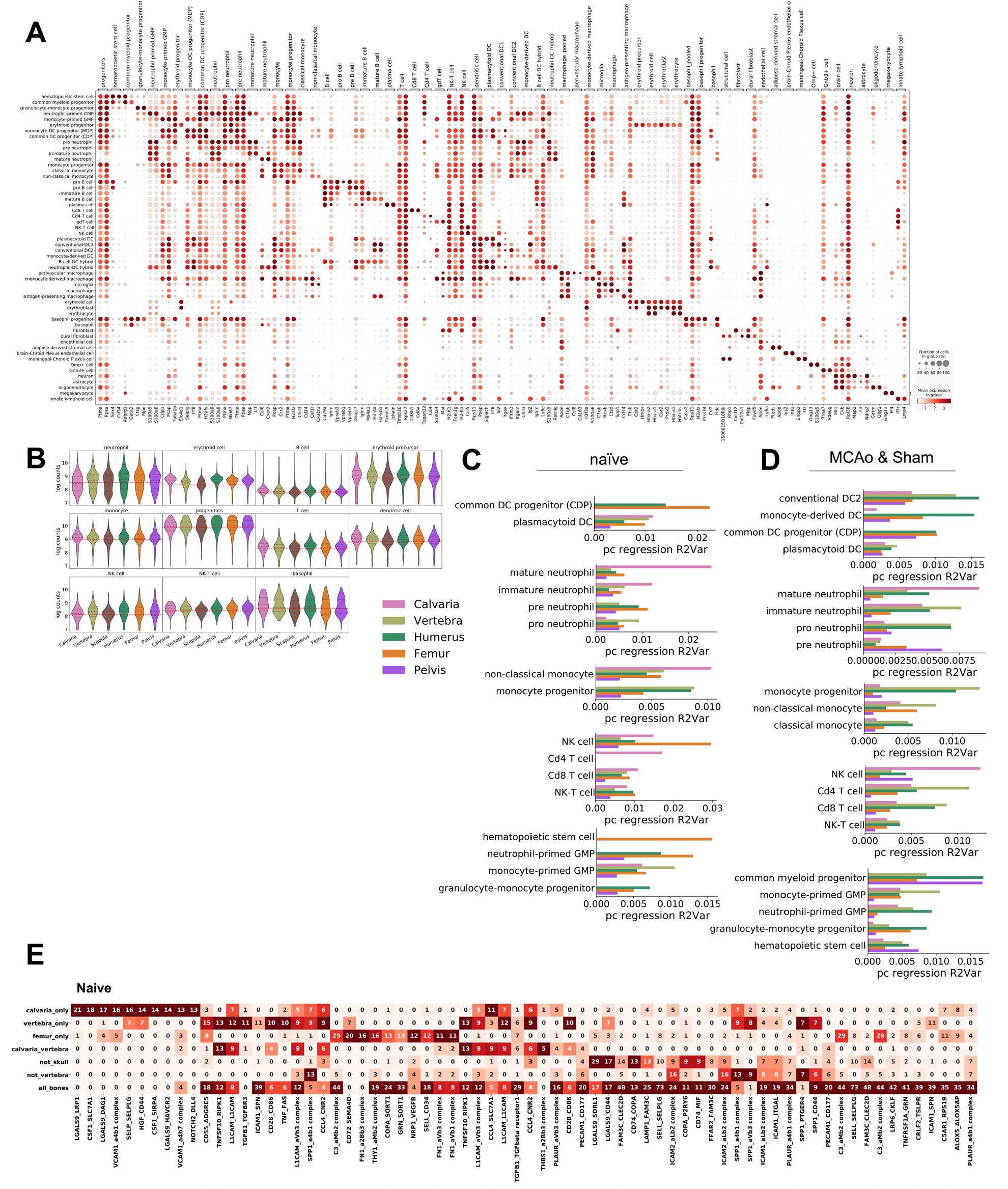
Marker genes for fine cell types and single cell RNA sequencing, QC measure, PC regression for fine cell-types and ligand-receptor pairs in naïve condition. (A) Dot plot shows coarse and fine annotated cell types and their marker genes. Cell types start with the coarse cell type annotation with one known marker gene from the literature and the highest scoring differentially expressed gene in our dataset. This is followed by fine cell types under this coarse cell type. Again one marker gene from the literature is shown, and additionally the highest scoring differentially expressed gene within the coarse cell type. (B) Log counts in bones over coarse cell types are shown. Scapula, among other bones shows less log counts in all cell types hinting a systemic error in scapula cells. (C) Divergence of each bone’s cell type population from the population of all other bones measured via variance explained based on PC regression for fine cell type annotations in naïve condition. (D) Divergence of each bone’s cell type population from the population of all other bones measured via variance explained based on PC regression for fine cell type annotations in MCAo and sham combined populations. (E) Number of cell type combinations for top ranking ligand-receptor pairs that are measured in the groups: only calvaria, only vertebra, only femur, calvaria & vertebra, all except calvaria, all except vertebra, and all bones. The top 10 pairs ranked by number of cell type pairs of each group are shown in naive condition.

**Fig. S3.**
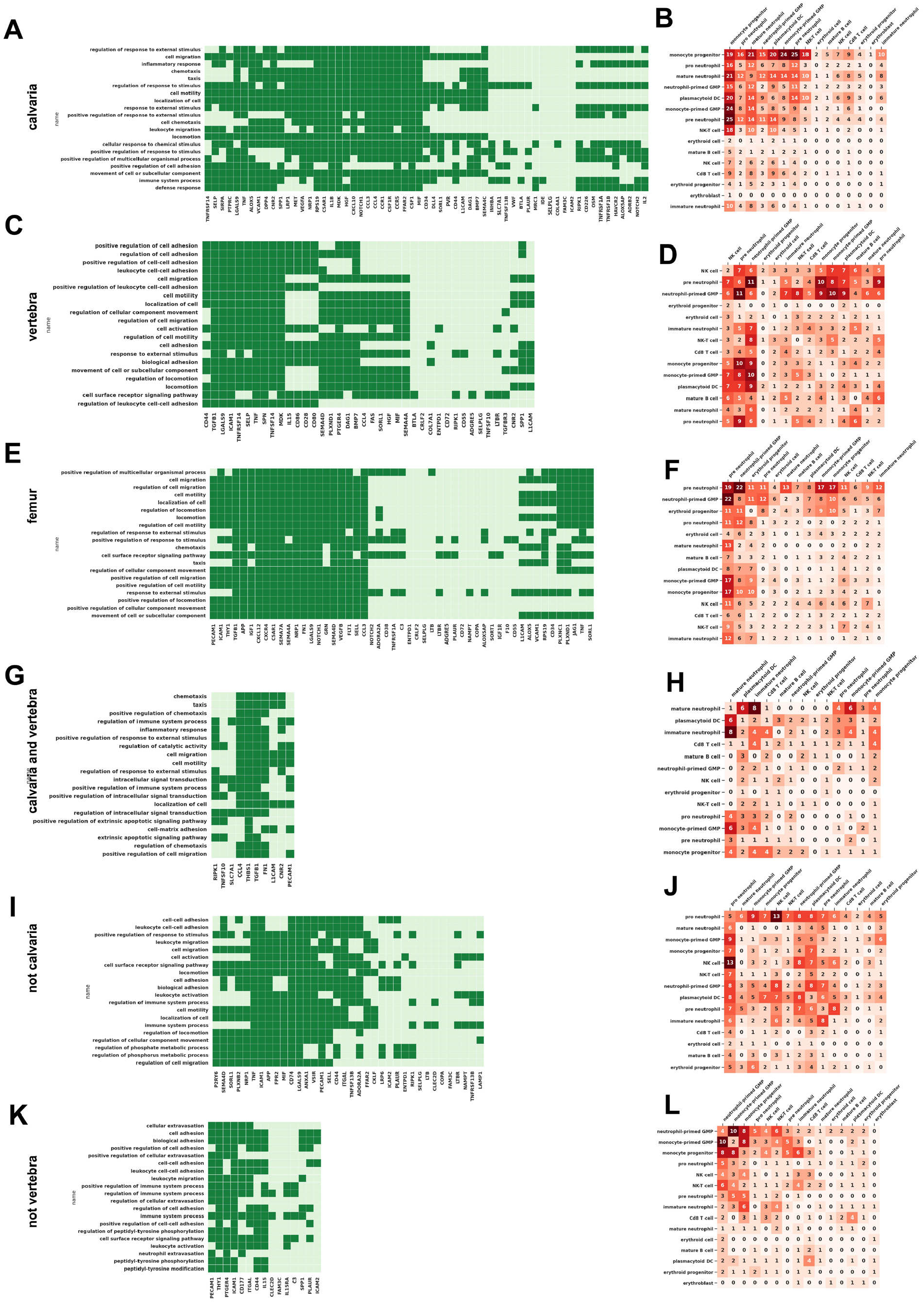
Unique interaction partners in bones in naïve condition. (A) Genes and their 20 most significant GO terms of ligand receptor pairs uniquely measured in calvaria. (B) Number of ligand receptor pairs uniquely measured in calvaria. (C) Genes and their 20 most significant GO terms of ligand receptor pairs uniquely measured in vertebra. (D) Number of ligand receptor pairs uniquely measured in vertebra. (E) Genes and their 20 most significant GO terms of ligand receptor pairs uniquely measured in femur. (F) Number of ligand receptor pairs uniquely measured in femur. (G) Genes and their 20 most significant GO terms of ligand receptor pairs uniquely measured in calvaria & vertebra. (H) Number of ligand receptor pairs uniquely measured in calvaria & vertebra. (I) Genes and their 20 most significant GO terms of ligand receptor pairs uniquely measured in all bones except for calvaria. (J) Number of ligand receptor pairs uniquely measured in all bones except for calvaria. (K) Genes and their 20 most significant GO terms of ligand receptor pairs uniquely measured in all bones except for vertebra. (L) Number of ligand receptor pairs uniquely measured in all bones except for vertebra. Genes were ordered by a linkage clustering based on their GO term similarity in (A, C, E, G, I, K).

**Fig. S4.**
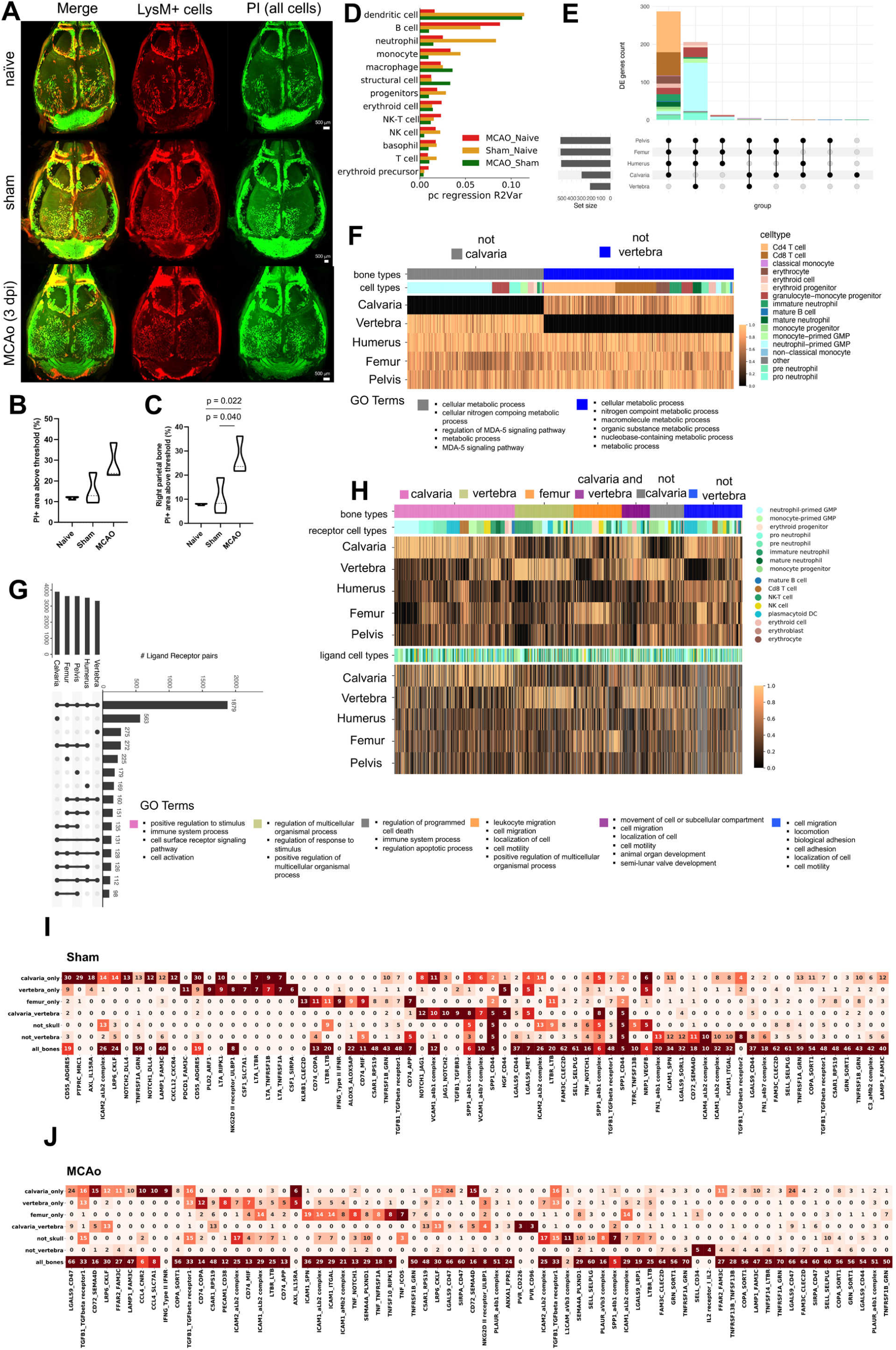
Structural and transcriptomic changes in injury conditions. (A) Whole head clearing of LysM mice in naïve, sham and MCAo (stroke was performed on the left side) condition show increased PI and LysM signal after MCAo. (B) Quantification of PI signal in the frontal and parietal bones show a strong trend [F(2,6) = 5.027, p = 0.522] for increased PI signal in MCAo condition compared to sham (p = 0.124) and naïve condition (p = 0.053). (C) Quantification of PI signal in the contralateral parietal skull bone of show increase [F(2,6) = 8.323, p = 0.019] in PI signal in MCAo condition compared to sham (p = 0.040) and naïve (p = 0.022) condition. (n=3 per group, C57BL6/J mice has been used for naïve condition quantification). dpi: days post injury. (D) Divergence of cell type populations between condition pairs measured via variance explained based on PC regression. (E) The upset plot demonstrates the common differentially expressed genes among bones and the cell types that these genes are higher expressed in, in sham condition. (F) The differentially expressed genes are represented with a mean expression heatmap with the corresponding GO terms, in sham condition. (G) The upset plot demonstrates the ligand-receptor pairs expressed in bones in sham conditions. (H) Ligand-receptor analysis in sham conditions are shown with the corresponding genes and GO terms in the heatmap in sham condition. The minimal mean expression among bones is set to zero and the maximal to one. (n=3 pooled animals for sham. “not calvaria/vertebra” means that, that module is present in all other bones except for ‘calvaria/vertebra’ in both heatmaps. (I) Number of cell type combinations for top ranking ligand-receptor pairs that are measured in the groups: only calvaria, only vertebra, only femur, calvaria & vertebra, all except calvaria, all except vertebra, and all bones. The top 10 pairs ranked by number of cell type pairs of each group are shown in sham condition. (J) Number of cell type combinations for top ranking ligand-receptor pairs that are measured in the groups: only calvaria, only vertebra, only femur, calvaria & vertebra, all except calvaria, all except vertebra, and all bones. The top 10 pairs ranked by number of cell type pairs of each group are shown in MCAo condition.

**Fig. S5.**
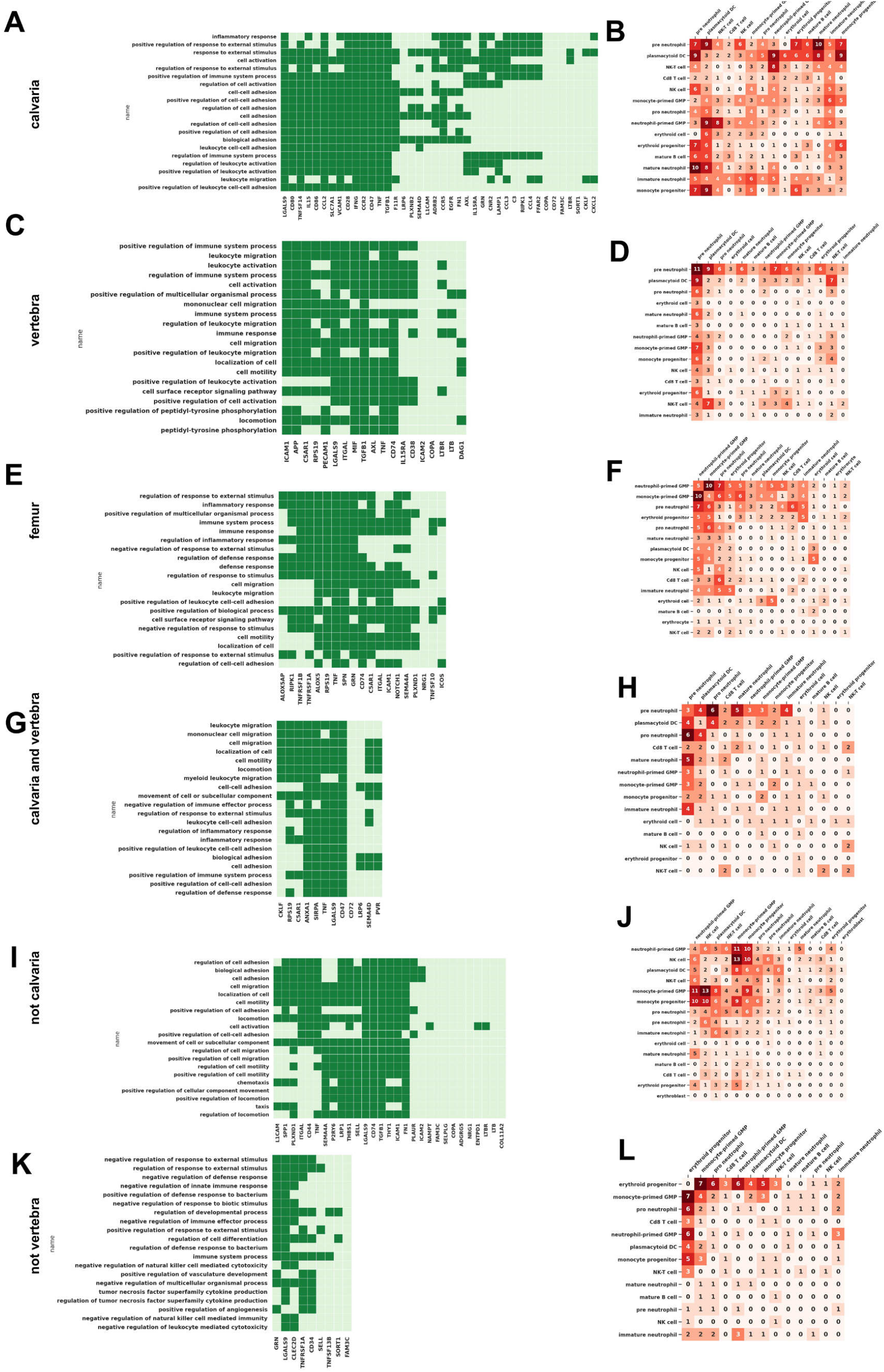
Unique interaction partners in bones in MCAo condition. (A) Genes and their 20 most significant GO terms of ligand receptor pairs uniquely measured in calvaria. (B) Number of ligand receptor pairs uniquely measured in calvaria. (C) Genes and their 20 most significant GO terms of ligand receptor pairs uniquely measured in vertebra. (D) Number of ligand receptor pairs uniquely measured in vertebra. (E) Genes and their 20 most significant GO terms of ligand receptor pairs uniquely measured in femur. (F) Number of ligand receptor pairs uniquely measured in femur. (G) Genes and their 20 most significant GO terms of ligand receptor pairs uniquely measured in calvaria & vertebra. (H) Number of ligand receptor pairs uniquely measured in calvaria & vertebra. (I) Genes and their 20 most significant GO terms of ligand receptor pairs uniquely measured in all bones except for calvaria. (J) Number of ligand receptor pairs uniquely measured in all bones except for calvaria. (K) Genes and their 20 most significant GO terms of ligand receptor pairs uniquely measured in all bones except for vertebra. (L) Number of ligand receptor pairs uniquely measured in all bones except for vertebra. Genes were ordered by a linkage clustering based on their GO term similarity in (A, C, E, G, I, K).

**Fig. S6.**
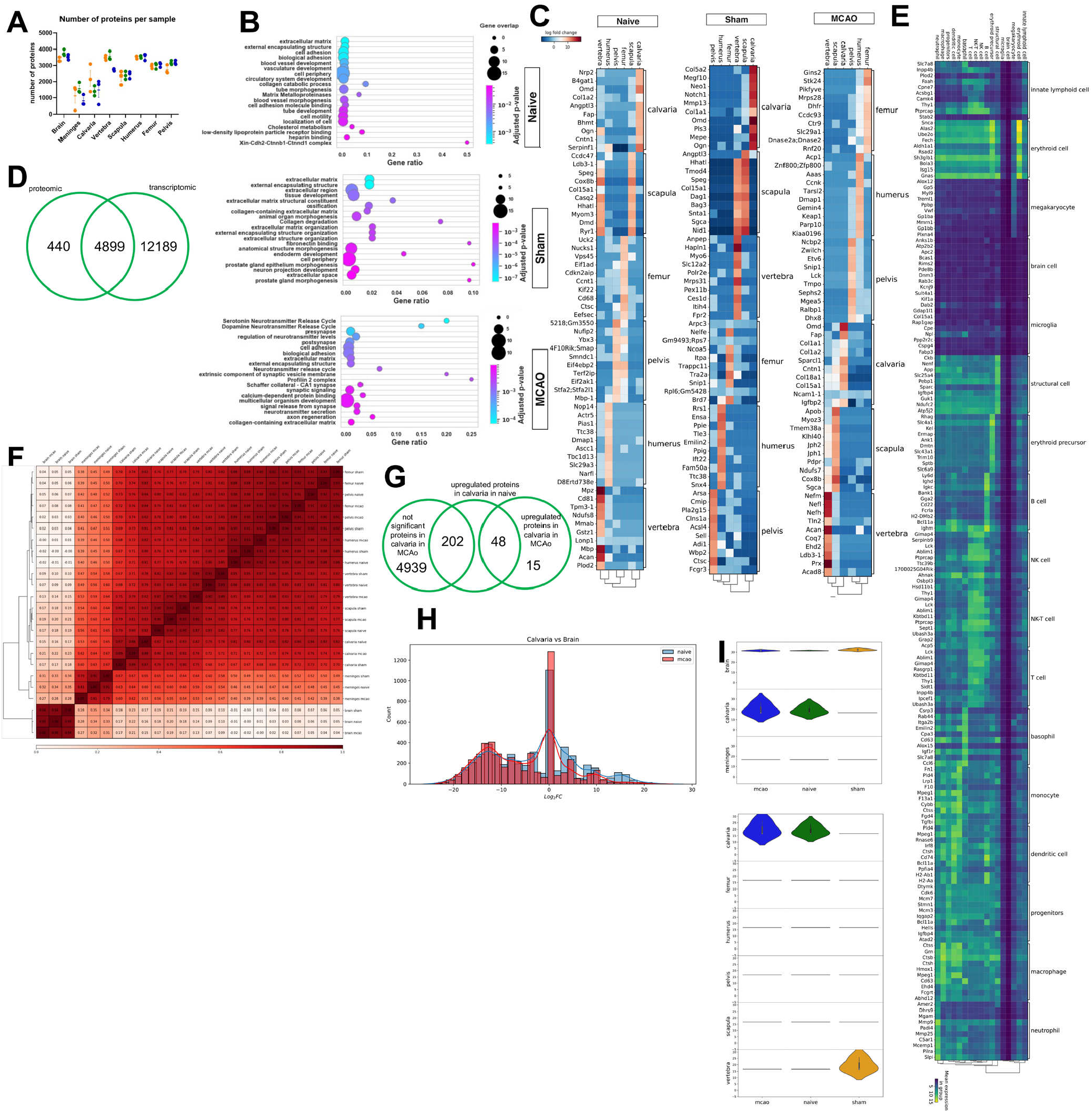
Calvaria shows cues of brain communication after MCAO. (A)Number of proteins per sample as a quality control measure. (B)GO terms from uniquely detected proteins in the calvaria in naïve, sham and MCAo condition. (C)Highly expressed markers in naïve, sham and MCAo condition are shown in a matrix plot. (D)Common genes detected in the proteomic and transcriptomic datasets shown in a Venn diagram. (E)Marker genes are shown for deconvoluted cell types. (F)Correlation matrix among all bones, meninges and brain in all conditions are shown using Pearson’s correlation with average linkage. (G)Venn diagram shows the changes in the volcano plot shown in Figure 4I. Plot depicts brain and calvaria samples becoming similar after the course of stroke. (H)Histogram shows changes in the log fold change of detected proteins in naïve and MCAo condition between the calvaria and the brain. (I)Violin plot demonstrates Gap43 expression in brain, meninges and calvaria (upper) and bones (lower) in three conditions.

**Fig. S7.**
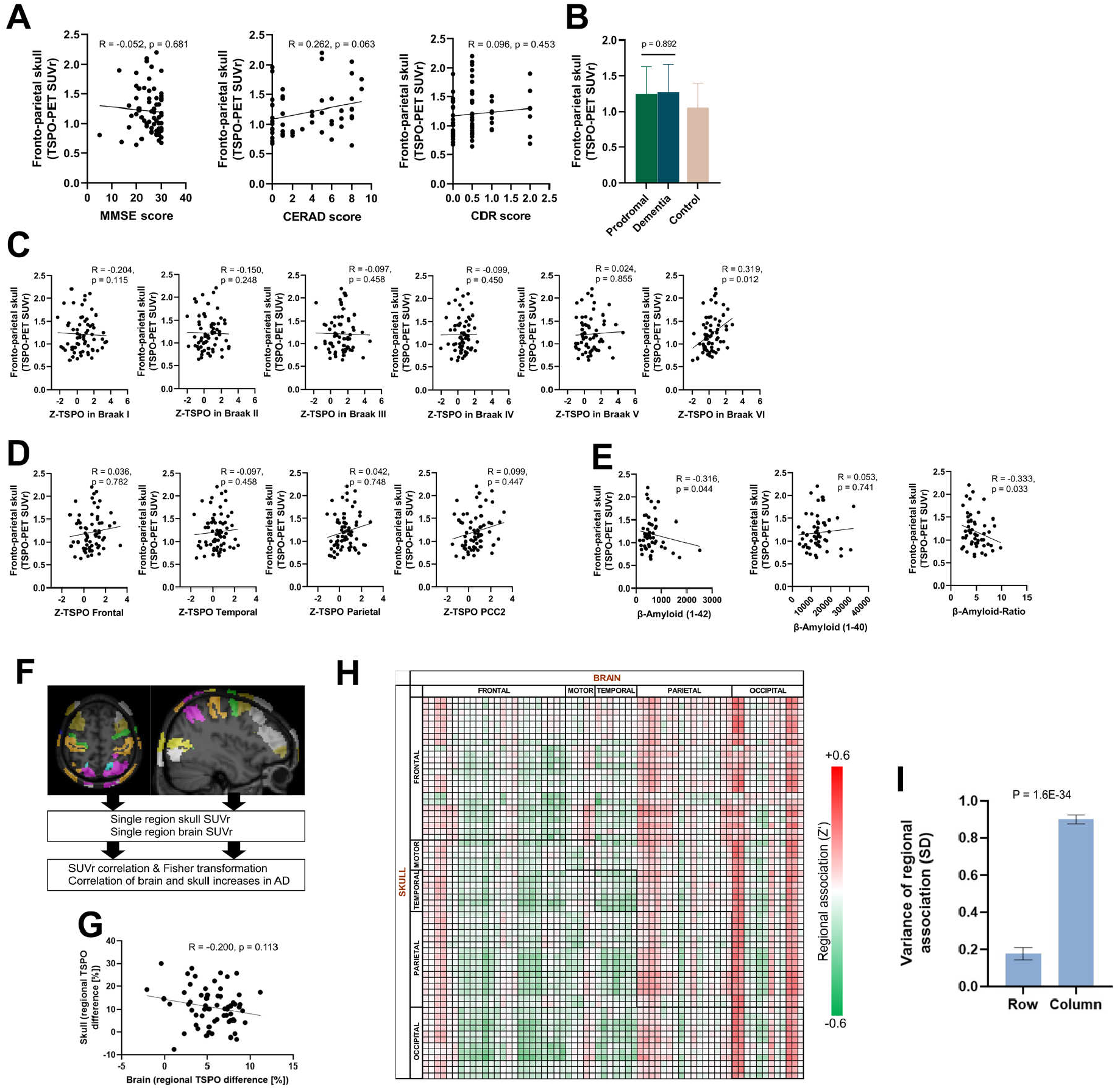
Human TSPO-PET scans reveal Aβ42 and higher Braak stage region associated TSPO signal in the calvaria. (A) Fronto-parietal skull TSPO signal from human AD patients show no significant correlation with clinical severity as demonstrated by MMSE (p=0.681), CERAD (p=0.063) and CDR (p=0.453) scorings respectively. (B) Fronto-parietal skull TSPO signal shows a homogenous increase in prodromal and dementia subgroups for Alzheimer’s disease compared to control patients (prodromal vs. dementia: p=0.892). (C) Fronto-parietal skull TSPO signal shows a positive association only with brain TSPO signal in the Braak VI stage region. (p=0.115 for Braak I, p=0.248 for Braak II, p=0.458 for Braak III, p=0.450 for Braak IV, p=0.855 for Braak V and p=0.012 for Braak VI). (D) Fronto-parietal skull TSPO signal is not significantly associated with brain TSPO signal in any β-amyloid related regions: frontal (p=0.782), temporal (p=0.458), parietal (p=0.748), and posterior cingulate cortex/ precuneus (p=0.447). (E) Fronto-parietal skull TSPO signal is correlated with β-amyloid42 (p=0.044) but not β-amyloid40 (p=0.741) in cerebrospinal fluid, also reflected by the significant negative correlation of the β-amyloid ratio (p=0.033). (F) The association between brain and corresponding skull regions were computed after ROI selection, SUVr correlation and Fisher transformation. (G) Correlation between regional TSPO differences in patients with AD vs. controls for the brain vs the skull. The correlation is not significant (p=0.113). (H) Correlation matrix showing TSPO-PET signal associations in brain regions vs skull regions. Parietal and occipital brain regions display high correlation with all skull regions. i, (I) The high correlation between single brain regions with all skull regions is further shown by row vs. column variances in the matrix plot in h (p=1.6E-34). The significant difference suggests that brain inflammation in any region is reflected to the whole skull rather than a corresponding region.

