## Supplementary material for "Multi-omics and 3D-imaging reveal bone heterogeneity and unique calvaria cells in neuroinflammation": Table S1

| level1_all | Brain | Meninges | Skull | Vertebra | Scapula | Humerus | Pelvis | Femur |
|---|---|---|---|---|---|---|---|---|
| progenitors | 25 | 128 | 466 | 786 | 734 | 1114 | 1030 | 1294 |
| neutrophil | 36 | 792 | 7608 | 7702 | 6796 | 8538 | 9237 | 10138 |
| monocyte | 14 | 103 | 865 | 1382 | 1355 | 1354 | 1591 | 1599 |
| B cell | 4 | 201 | 3393 | 2358 | 2937 | 3192 | 3204 | 3322 |
| T cell | 4 | 165 | 622 | 347 | 382 | 390 | 487 | 546 |
| NK cell | 0 | 65 | 255 | 162 | 226 | 130 | 164 | 154 |
| NK-T cell | 0 | 85 | 128 | 126 | 146 | 168 | 157 | 199 |
| dendritic cell | 18 | 375 | 190 | 235 | 334 | 344 | 333 | 374 |
| macrophage | 90 | 2302 | 8 | 20 | 28 | 22 | 18 | 14 |
| microglia | 654 | 57 | 0 | 1 | 1 | 1 | 1 | 0 |
| erythroid precursor | 42 | 152 | 1040 | 1717 | 1968 | 3055 | 2931 | 4199 |
| erythroid cell | 22 | 907 | 5152 | 3785 | 3945 | 5094 | 5546 | 7154 |
| basophil | 0 | 3 | 60 | 88 | 75 | 113 | 141 | 129 |
| structural cell | 187 | 3808 | 34 | 54 | 91 | 54 | 51 | 59 |
| brain cell | 1724 | 29 | 0 | 0 | 0 | 0 | 2 | 0 |
| megakaryocyte | 1 | 1 | 2 | 1 | 2 | 6 | 0 | 6 |
| innate lymphoid cell | 1 | 51 | 6 | 6 | 7 | 8 | 8 | 4 |

| level1_naive | Brain | Meninges | Skull | Vertebra | Scapula | Humerus | Pelvis | Femur |
|---|---|---|---|---|---|---|---|---|
| progenitors | 5 | 46 | 109 | 125 | 226 | 226 | 196 | 256 |
| neutrophil | 6 | 75 | 1635 | 1040 | 1906 | 1708 | 1862 | 2020 |
| monocyte | 3 | 13 | 147 | 185 | 461 | 262 | 318 | 295 |
| B cell | 0 | 42 | 712 | 407 | 991 | 938 | 790 | 858 |
| T cell | 1 | 22 | 74 | 35 | 65 | 47 | 52 | 63 |
| NK cell | 0 | 7 | 52 | 26 | 75 | 30 | 30 | 34 |
| NK-T cell | 0 | 13 | 35 | 21 | 54 | 40 | 44 | 43 |
| dendritic cell | 6 | 64 | 51 | 50 | 135 | 82 | 76 | 89 |
| macrophage | 16 | 305 | 3 | 9 | 9 | 8 | 7 | 2 |
| microglia | 86 | 19 | 0 | 0 | 0 | 0 | 0 | 0 |
| erythroid precursor | 8 | 39 | 396 | 448 | 987 | 1243 | 1131 | 1581 |
| erythroid cell | 6 | 210 | 1719 | 1075 | 1795 | 1921 | 1997 | 2487 |
| basophil | 0 | 2 | 13 | 15 | 24 | 28 | 48 | 34 |
| structural cell | 75 | 1253 | 8 | 12 | 18 | 10 | 13 | 6 |
| brain cell | 294 | 9 | 0 | 0 | 0 | 0 | 0 | 0 |
| megakaryocyte | 0 | 1 | 0 | 0 | 0 | 1 | 0 | 3 |
| innate lymphoid cell | 0 | 6 | 1 | 0 | 5 | 1 | 3 | 0 |

| level1_sham | Brain | Meninges | Skull | Vertebra | Scapula | Humerus | Pelvis | Femur |
|---|---|---|---|---|---|---|---|---|
| progenitors | 0 | 32 | 75 | 69 | 203 | 216 | 279 | 292 |
| neutrophil | 3 | 48 | 1492 | 679 | 1429 | 1603 | 2198 | 2196 |
| monocyte | 1 | 5 | 163 | 96 | 228 | 234 | 354 | 297 |
| B cell | 0 | 11 | 838 | 238 | 562 | 560 | 761 | 578 |
| T cell | 0 | 14 | 193 | 27 | 96 | 102 | 142 | 120 |
| NK cell | 0 | 0 | 59 | 21 | 48 | 33 | 43 | 37 |
| NK-T cell | 0 | 9 | 28 | 17 | 27 | 27 | 31 | 39 |
| dendritic cell | 1 | 29 | 55 | 27 | 83 | 98 | 123 | 104 |
| macrophage | 13 | 217 | 2 | 3 | 4 | 5 | 10 | 5 |
| microglia | 116 | 10 | 0 | 1 | 0 | 0 | 1 | 0 |
| erythroid precursor | 3 | 45 | 237 | 249 | 369 | 633 | 736 | 857 |
| erythroid cell | 0 | 93 | 1254 | 388 | 777 | 1096 | 1414 | 1472 |
| basophil | 0 | 0 | 11 | 9 | 14 | 18 | 37 | 25 |
| structural cell | 28 | 714 | 9 | 17 | 13 | 15 | 29 | 23 |
| brain cell | 340 | 11 | 0 | 0 | 0 | 0 | 2 | 0 |
| megakaryocyte | 0 | 0 | 1 | 0 | 0 | 1 | 0 | 0 |
| innate lymphoid cell | 0 | 8 | 0 | 0 | 0 | 1 | 2 | 2 |

| level1_MCAo | Brain | Meninges | Skull | Vertebra | Scapula | Humerus | Pelvis | Femur |
|---|---|---|---|---|---|---|---|---|
| progenitors | 20 | 50 | 282 | 592 | 305 | 672 | 555 | 746 |
| neutrophil | 27 | 669 | 4481 | 5983 | 3461 | 5227 | 5177 | 5922 |
| monocyte | 10 | 85 | 555 | 1101 | 666 | 858 | 919 | 1007 |
| B cell | 4 | 148 | 1843 | 1713 | 1384 | 1694 | 1653 | 1886 |
| T cell | 3 | 129 | 355 | 285 | 221 | 241 | 293 | 363 |
| NK cell | 0 | 58 | 144 | 115 | 103 | 67 | 91 | 83 |
| NK-T cell | 0 | 63 | 65 | 88 | 65 | 101 | 82 | 117 |
| dendritic cell | 11 | 282 | 84 | 158 | 116 | 164 | 134 | 181 |
| macrophage | 61 | 1780 | 3 | 8 | 15 | 9 | 1 | 7 |
| microglia | 452 | 28 | 0 | 0 | 1 | 1 | 0 | 0 |
| erythroid precursor | 31 | 68 | 407 | 1020 | 612 | 1179 | 1064 | 1761 |
| erythroid cell | 16 | 604 | 2179 | 2322 | 1373 | 2077 | 2135 | 3195 |
| basophil | 0 | 1 | 36 | 64 | 37 | 67 | 56 | 70 |
| structural cell | 84 | 1841 | 17 | 25 | 60 | 29 | 9 | 30 |
| brain cell | 1090 | 9 | 0 | 0 | 0 | 0 | 0 | 0 |
| megakaryocyte | 1 | 0 | 1 | 1 | 2 | 4 | 0 | 3 |
| innate lymphoid cell | 1 | 37 | 5 | 6 | 2 | 6 | 3 | 2 |

| level2_all | Brain | Meninges | Skull | Vertebra | Scapula | Humerus | Pelvis | Femur |
|---|---|---|---|---|---|---|---|---|
| hematopoietic stem cell | 0 | 5 | 42 | 56 | 64 | 101 | 84 | 112 |
| common myeloid progenitor | 0 | 4 | 26 | 45 | 37 | 53 | 44 | 63 |
| granulocyte-monocyte progenitor | 24 | 95 | 83 | 149 | 125 | 259 | 206 | 366 |
| neutrophil-primed GMP | 1 | 9 | 102 | 218 | 215 | 298 | 263 | 276 |
| monocyte-primed GMP | 0 | 9 | 193 | 302 | 264 | 373 | 398 | 453 |
| erythroid progenitor | 40 | 124 | 566 | 899 | 1119 | 1750 | 1613 | 2414 |
| monocyte-DC progenitor (MDP) | 0 | 6 | 20 | 16 | 29 | 30 | 35 | 24 |
| common DC progenitor (CDP) | 0 | 0 | 24 | 23 | 29 | 54 | 37 | 52 |
| pro neutrophil | 0 | 26 | 167 | 422 | 409 | 716 | 607 | 658 |
| pre neutrophil | 19 | 66 | 125 | 221 | 164 | 230 | 281 | 384 |
| immature neutrophil | 3 | 147 | 2160 | 2954 | 2529 | 3379 | 3674 | 3960 |
| mature neutrophil | 14 | 553 | 5156 | 4105 | 3694 | 4213 | 4675 | 5136 |
| monocyte progenitor | 0 | 9 | 151 | 246 | 190 | 271 | 329 | 315 |
| classical monocyte | 14 | 54 | 497 | 888 | 647 | 775 | 901 | 937 |
| non-classical monocyte | 0 | 40 | 217 | 248 | 518 | 308 | 361 | 347 |
| pro B cell | 0 | 8 | 122 | 65 | 124 | 194 | 123 | 175 |
| pre B cell | 0 | 18 | 188 | 121 | 253 | 239 | 157 | 194 |
| immature B cell | 0 | 25 | 159 | 141 | 253 | 223 | 232 | 200 |
| mature B cell | 4 | 147 | 2916 | 2024 | 2303 | 2514 | 2675 | 2731 |
| plasma cell | 0 | 3 | 8 | 7 | 4 | 22 | 17 | 22 |
| Cd8 T cell | 3 | 39 | 376 | 225 | 235 | 265 | 328 | 371 |
| Cd4 T cell | 0 | 42 | 238 | 118 | 144 | 120 | 149 | 168 |
| gdT cell | 1 | 84 | 8 | 4 | 3 | 5 | 10 | 7 |
| NK-T cell | 0 | 85 | 128 | 126 | 146 | 168 | 157 | 199 |
| NK cell | 0 | 65 | 255 | 162 | 226 | 130 | 164 | 154 |
| plasmacytoid DC | 0 | 18 | 95 | 129 | 157 | 169 | 160 | 191 |
| conventional DC1 | 1 | 49 | 16 | 7 | 14 | 13 | 15 | 5 |
| conventional DC2 | 0 | 149 | 30 | 52 | 65 | 44 | 35 | 48 |
| monocyte-derived DC | 17 | 158 | 22 | 16 | 67 | 57 | 81 | 71 |
| B cell-DC hybrid | 0 | 0 | 2 | 7 | 1 | 5 | 4 | 2 |
| neutrophil-DC hybrid | 0 | 1 | 1 | 1 | 1 | 2 | 1 | 5 |
| perivascular macrophage | 34 | 1416 | 0 | 0 | 5 | 0 | 3 | 3 |
| monocyte-derived macrophage | 15 | 280 | 1 | 0 | 0 | 0 | 0 | 0 |

| | | | | | | | | |
|---|---|---|---|---|---|---|---|---|
| **microglia** | 654 | 57 | 0 | 1 | 1 | 1 | 1 | 0 |
| **macrophage** | 34 | 29 | 5 | 6 | 5 | 2 | 7 | 4 |
| **antigen-presenting macrophage** | 7 | 577 | 2 | 14 | 18 | 20 | 8 | 7 |
| **erythroid cell** | 2 | 28 | 474 | 818 | 849 | 1305 | 1318 | 1785 |
| **erythroblast** | 3 | 167 | 1944 | 1984 | 1937 | 2785 | 2623 | 3790 |
| **erythrocyte** | 19 | 740 | 3208 | 1801 | 2008 | 2309 | 2923 | 3364 |
| **basophil progenitor** | 0 | 0 | 6 | 10 | 8 | 13 | 18 | 10 |
| **basophil** | 0 | 3 | 54 | 78 | 67 | 100 | 123 | 119 |
| **fibroblast** | 0 | 5 | 7 | 7 | 17 | 34 | 10 | 31 |
| **dural fibroblast** | 30 | 1591 | 4 | 19 | 8 | 6 | 10 | 3 |
| **endothelial cell** | 33 | 55 | 2 | 1 | 4 | 0 | 2 | 2 |
| **adipose-derived stromal cell** | 5 | 31 | 0 | 0 | 1 | 0 | 0 | 0 |
| **brain-Chroid Plexus endothelial cell** | 17 | 0 | 0 | 0 | 0 | 0 | 0 | 0 |
| **meningeal-Choroid Plexus cell** | 46 | 1724 | 5 | 6 | 25 | 2 | 10 | 5 |
| **Omp+ cell** | 0 | 159 | 0 | 0 | 0 | 0 | 0 | 0 |
| **Gnb3+ cell** | 56 | 243 | 16 | 21 | 36 | 12 | 19 | 18 |
| **neuron** | 373 | 1 | 0 | 0 | 0 | 0 | 0 | 0 |
| **astrocyte** | 1187 | 24 | 0 | 0 | 0 | 0 | 1 | 0 |
| **oligodendrocyte** | 164 | 4 | 0 | 0 | 0 | 0 | 1 | 0 |
| **megakaryocyte** | 1 | 1 | 2 | 1 | 2 | 6 | 0 | 6 |
| **innate lymphoid cell** | 1 | 51 | 6 | 6 | 7 | 8 | 8 | 4 |

| level2_naive | Brain | Meninges | Skull | Vertebra | Scapula | Humerus | Pelvis | Femur |
|---|---|---|---|---|---|---|---|---|
| hematopoietic stem cell | 0 | 1 | 16 | 5 | 19 | 17 | 18 | 31 |
| common myeloid progenitor | 0 | 0 | 10 | 6 | 12 | 13 | 11 | 19 |
| granulocyte-monocyte progenitor | 5 | 42 | 9 | 20 | 38 | 47 | 31 | 65 |
| neutrophil-primed GMP | 0 | 1 | 11 | 17 | 43 | 34 | 25 | 27 |
| monocyte-primed GMP | 0 | 2 | 54 | 72 | 95 | 102 | 92 | 101 |
| erythroid progenitor | 8 | 32 | 197 | 238 | 549 | 703 | 640 | 952 |
| monocyte-DC progenitor (MDP) | 0 | 0 | 9 | 5 | 19 | 13 | 19 | 13 |
| common DC progenitor (CDP) | 0 | 0 | 13 | 5 | 16 | 23 | 15 | 22 |
| pro neutrophil | 0 | 1 | 21 | 56 | 87 | 103 | 85 | 101 |
| pre neutrophil | 4 | 28 | 16 | 22 | 68 | 38 | 45 | 48 |
| immature neutrophil | 0 | 21 | 419 | 375 | 561 | 552 | 595 | 702 |
| mature neutrophil | 2 | 25 | 1179 | 587 | 1190 | 1015 | 1137 | 1169 |
| monocyte progenitor | 0 | 0 | 18 | 31 | 51 | 51 | 63 | 58 |
| classical monocyte | 3 | 3 | 6 | 2 | 17 | 4 | 7 | 6 |
| non-classical monocyte | 0 | 10 | 123 | 152 | 393 | 207 | 248 | 231 |
| pro B cell | 0 | 8 | 117 | 54 | 120 | 188 | 119 | 165 |
| pre B cell | 0 | 9 | 179 | 107 | 244 | 227 | 147 | 183 |
| immature B cell | 0 | 15 | 58 | 57 | 151 | 112 | 120 | 96 |
| mature B cell | 0 | 10 | 358 | 189 | 473 | 408 | 398 | 409 |
| plasma cell | 0 | 0 | 0 | 0 | 3 | 3 | 6 | 5 |
| Cd8 T cell | 1 | 6 | 48 | 26 | 44 | 30 | 39 | 48 |
| Cd4 T cell | 0 | 4 | 25 | 7 | 20 | 16 | 13 | 11 |
| gdT cell | 0 | 12 | 1 | 2 | 1 | 1 | 0 | 4 |
| NK-T cell | 0 | 13 | 35 | 21 | 54 | 40 | 44 | 43 |
| NK cell | 0 | 7 | 52 | 26 | 75 | 30 | 30 | 34 |
| plasmacytoid DC | 0 | 5 | 25 | 25 | 67 | 41 | 40 | 52 |
| conventional DC1 | 0 | 6 | 2 | 0 | 4 | 4 | 8 | 1 |
| conventional DC2 | 0 | 16 | 8 | 12 | 40 | 11 | 12 | 13 |
| monocyte-derived DC | 6 | 37 | 1 | 6 | 6 | 0 | 1 | 0 |
| B cell-DC hybrid | 0 | 0 | 1 | 2 | 1 | 2 | 0 | 0 |
| neutrophil-DC hybrid | 0 | 0 | 1 | 0 | 1 | 1 | 0 | 1 |
| perivascular macrophage | 9 | 192 | 0 | 0 | 1 | 0 | 0 | 0 |
| microglia | 86 | 19 | 0 | 0 | 0 | 0 | 0 | 0 |

| | | | | | | | | |
|---|---|---|---|---|---|---|---|---|
| **macrophage** | 6 | 8 | 2 | 2 | 2 | 0 | 1 | 0 |
| **antigen-presenting macrophage** | 1 | 105 | 1 | 7 | 6 | 8 | 6 | 2 |
| **erythroid cell** | 0 | 7 | 199 | 210 | 438 | 540 | 491 | 629 |
| **erythroblast** | 0 | 46 | 804 | 612 | 910 | 1202 | 1056 | 1496 |
| **erythrocyte** | 6 | 164 | 915 | 463 | 885 | 719 | 941 | 991 |
| **basophil progenitor** | 0 | 0 | 2 | 6 | 5 | 7 | 9 | 1 |
| **basophil** | 0 | 2 | 11 | 9 | 19 | 21 | 39 | 33 |
| **fibroblast** | 0 | 1 | 2 | 1 | 7 | 6 | 3 | 5 |
| **dural fibroblast** | 10 | 369 | 2 | 3 | 2 | 1 | 1 | 0 |
| **endothelial cell** | 10 | 16 | 0 | 0 | 1 | 0 | 1 | 0 |
| **adipose-derived stromal cell** | 2 | 7 | 0 | 0 | 0 | 0 | 0 | 0 |
| **brain-Chroid Plexus endothelial cell** | 17 | 0 | 0 | 0 | 0 | 0 | 0 | 0 |
| **meningeal-Choroid Plexus cell** | 18 | 685 | 0 | 2 | 2 | 0 | 2 | 0 |
| **Omp+ cell** | 0 | 102 | 0 | 0 | 0 | 0 | 0 | 0 |
| **Gnb3+ cell** | 18 | 73 | 4 | 6 | 6 | 3 | 6 | 1 |
| **neuron** | 43 | 1 | 0 | 0 | 0 | 0 | 0 | 0 |
| **astrocyte** | 230 | 7 | 0 | 0 | 0 | 0 | 0 | 0 |
| **oligodendrocyte** | 21 | 1 | 0 | 0 | 0 | 0 | 0 | 0 |
| **megakaryocyte** | 0 | 1 | 0 | 0 | 0 | 1 | 0 | 3 |
| **innate lymphoid cell** | 0 | 6 | 1 | 0 | 5 | 1 | 3 | 0 |

| level2_sham | Brain | Meninges | Skull | Vertebra | Scapula | Humerus | Pelvis | Femur |
|---|---|---|---|---|---|---|---|---|
| hematopoietic stem cell | 0 | 1 | 4 | 5 | 14 | 22 | 25 | 19 |
| common myeloid progenitor | 0 | 0 | 7 | 6 | 10 | 12 | 7 | 12 |
| granulocyte-monocyte progenitor | 0 | 28 | 14 | 12 | 41 | 49 | 57 | 75 |
| neutrophil-primed GMP | 0 | 1 | 17 | 22 | 71 | 63 | 89 | 75 |
| monocyte-primed GMP | 0 | 2 | 31 | 22 | 64 | 67 | 101 | 105 |
| erythroid progenitor | 3 | 40 | 138 | 124 | 208 | 350 | 379 | 467 |
| monocyte-DC progenitor (MDP) | 0 | 0 | 2 | 2 | 3 | 3 | 0 | 6 |
| common DC progenitor (CDP) | 0 | 0 | 4 | 2 | 6 | 6 | 4 | 6 |
| pro neutrophil | 0 | 1 | 29 | 52 | 105 | 157 | 168 | 159 |
| pre neutrophil | 2 | 15 | 17 | 17 | 42 | 54 | 50 | 99 |
| immature neutrophil | 0 | 11 | 523 | 337 | 629 | 793 | 1089 | 1107 |
| mature neutrophil | 1 | 21 | 923 | 273 | 653 | 599 | 891 | 831 |
| monocyte progenitor | 0 | 0 | 30 | 17 | 51 | 64 | 83 | 65 |
| classical monocyte | 1 | 3 | 101 | 70 | 147 | 147 | 237 | 200 |
| non-classical monocyte | 0 | 2 | 32 | 9 | 30 | 23 | 34 | 32 |
| pro B cell | 0 | 0 | 1 | 0 | 0 | 0 | 1 | 1 |
| pre B cell | 0 | 0 | 0 | 1 | 1 | 2 | 1 | 1 |
| immature B cell | 0 | 0 | 43 | 9 | 15 | 24 | 37 | 19 |
| mature B cell | 0 | 11 | 794 | 228 | 546 | 531 | 722 | 555 |
| plasma cell | 0 | 0 | 0 | 0 | 0 | 3 | 0 | 2 |
| Cd8 T cell | 0 | 5 | 120 | 20 | 54 | 74 | 99 | 76 |
| Cd4 T cell | 0 | 5 | 70 | 7 | 41 | 28 | 38 | 43 |
| gdT cell | 0 | 4 | 3 | 0 | 1 | 0 | 5 | 1 |
| NK-T cell | 0 | 9 | 28 | 17 | 27 | 27 | 31 | 39 |
| NK cell | 0 | 0 | 59 | 21 | 48 | 33 | 43 | 37 |
| plasmacytoid DC | 0 | 0 | 23 | 17 | 19 | 27 | 36 | 24 |
| conventional DC1 | 0 | 3 | 5 | 0 | 2 | 2 | 3 | 2 |
| conventional DC2 | 0 | 8 | 6 | 3 | 3 | 9 | 4 | 6 |
| monocyte-derived DC | 1 | 18 | 17 | 2 | 53 | 52 | 75 | 66 |
| B cell-DC hybrid | 0 | 0 | 0 | 3 | 0 | 2 | 1 | 0 |
| perivascular macrophage | 10 | 159 | 0 | 0 | 0 | 0 | 2 | 1 |
| monocyte-derived macrophage | 0 | 3 | 0 | 0 | 0 | 0 | 0 | 0 |
| microglia | 116 | 10 | 0 | 1 | 0 | 0 | 1 | 0 |

| | | | | | | | | |
|---|---|---|---|---|---|---|---|---|
| **macrophage** | 3 | 10 | 2 | 3 | 2 | 1 | 6 | 2 |
| **antigen-presenting macrophage** | 0 | 45 | 0 | 0 | 2 | 4 | 2 | 2 |
| **erythroid cell** | 0 | 5 | 99 | 125 | 161 | 283 | 357 | 390 |
| **erythroblast** | 0 | 23 | 449 | 291 | 421 | 702 | 709 | 867 |
| **erythrocyte** | 0 | 70 | 805 | 97 | 356 | 394 | 705 | 605 |
| **basophil progenitor** | 0 | 0 | 2 | 0 | 1 | 0 | 4 | 3 |
| **basophil** | 0 | 0 | 9 | 9 | 13 | 18 | 33 | 22 |
| **fibroblast** | 0 | 0 | 2 | 2 | 1 | 6 | 3 | 8 |
| **dural fibroblast** | 4 | 305 | 1 | 2 | 1 | 2 | 7 | 1 |
| **endothelial cell** | 7 | 24 | 1 | 1 | 1 | 0 | 1 | 1 |
| **adipose-derived stromal cell** | 3 | 12 | 0 | 0 | 0 | 0 | 0 | 0 |
| **meningeal-Choroid Plexus cell** | 4 | 318 | 2 | 2 | 2 | 2 | 8 | 3 |
| **Omp+ cell** | 0 | 4 | 0 | 0 | 0 | 0 | 0 | 0 |
| **Gnb3+ cell** | 10 | 51 | 3 | 10 | 8 | 5 | 10 | 10 |
| **neuron** | 91 | 0 | 0 | 0 | 0 | 0 | 0 | 0 |
| **astrocyte** | 216 | 9 | 0 | 0 | 0 | 0 | 1 | 0 |
| **oligodendrocyte** | 33 | 2 | 0 | 0 | 0 | 0 | 1 | 0 |
| **megakaryocyte** | 0 | 0 | 1 | 0 | 0 | 1 | 0 | 0 |
| **innate lymphoid cell** | 0 | 8 | 0 | 0 | 0 | 1 | 2 | 2 |

| level2_MCAo | Brain | Meninges | Skull | Vertebra | Scapula | Humerus | Pelvis | Femur |
|---|---|---|---|---|---|---|---|---|
| hematopoietic stem cell | 0 | 3 | 22 | 46 | 31 | 62 | 41 | 62 |
| common myeloid progenitor | 0 | 4 | 9 | 33 | 15 | 28 | 26 | 32 |
| granulocyte-monocyte progenitor | 19 | 25 | 60 | 117 | 46 | 163 | 118 | 226 |
| neutrophil-primed GMP | 1 | 7 | 74 | 179 | 101 | 201 | 149 | 174 |
| monocyte-primed GMP | 0 | 5 | 108 | 208 | 105 | 204 | 205 | 247 |
| erythroid progenitor | 29 | 52 | 231 | 537 | 362 | 697 | 594 | 995 |
| monocyte-DC progenitor (MDP) | 0 | 6 | 9 | 9 | 7 | 14 | 16 | 5 |
| common DC progenitor (CDP) | 0 | 0 | 7 | 16 | 7 | 25 | 18 | 24 |
| pro neutrophil | 0 | 24 | 117 | 314 | 217 | 456 | 354 | 398 |
| pre neutrophil | 13 | 23 | 92 | 182 | 54 | 138 | 186 | 237 |
| immature neutrophil | 3 | 115 | 1218 | 2242 | 1339 | 2034 | 1990 | 2151 |
| mature neutrophil | 11 | 507 | 3054 | 3245 | 1851 | 2599 | 2647 | 3136 |
| monocyte progenitor | 0 | 9 | 103 | 198 | 88 | 156 | 183 | 192 |
| classical monocyte | 10 | 48 | 390 | 816 | 483 | 624 | 657 | 731 |
| non-classical monocyte | 0 | 28 | 62 | 87 | 95 | 78 | 79 | 84 |
| pro B cell | 0 | 0 | 4 | 11 | 4 | 6 | 3 | 9 |
| pre B cell | 0 | 9 | 9 | 13 | 8 | 10 | 9 | 10 |
| immature B cell | 0 | 10 | 58 | 75 | 87 | 87 | 75 | 85 |
| mature B cell | 4 | 126 | 1764 | 1607 | 1284 | 1575 | 1555 | 1767 |
| plasma cell | 0 | 3 | 8 | 7 | 1 | 16 | 11 | 15 |
| Cd8 T cell | 2 | 28 | 208 | 179 | 137 | 161 | 190 | 247 |
| Cd4 T cell | 0 | 33 | 143 | 104 | 83 | 76 | 98 | 114 |
| gdT cell | 1 | 68 | 4 | 2 | 1 | 4 | 5 | 2 |
| NK-T cell | 0 | 63 | 65 | 88 | 65 | 101 | 82 | 117 |
| NK cell | 0 | 58 | 144 | 115 | 103 | 67 | 91 | 83 |
| plasmacytoid DC | 0 | 13 | 47 | 87 | 71 | 101 | 84 | 115 |
| conventional DC1 | 1 | 40 | 9 | 7 | 8 | 7 | 4 | 2 |
| conventional DC2 | 0 | 125 | 16 | 37 | 22 | 24 | 19 | 29 |
| monocyte-derived DC | 10 | 103 | 4 | 8 | 8 | 5 | 5 | 5 |
| B cell-DC hybrid | 0 | 0 | 1 | 2 | 0 | 1 | 3 | 2 |
| neutrophil-DC hybrid | 0 | 1 | 0 | 1 | 0 | 1 | 1 | 4 |
| perivascular macrophage | 15 | 1065 | 0 | 0 | 4 | 0 | 1 | 2 |
| monocyte-derived macrophage | 15 | 277 | 1 | 0 | 0 | 0 | 0 | 0 |

| | | | | | | | | |
|---|---|---|---|---|---|---|---|---|
| **microglia** | 452 | 28 | 0 | 0 | 1 | 1 | 0 | 0 |
| **macrophage** | 25 | 11 | 1 | 1 | 1 | 1 | 0 | 2 |
| **antigen-presenting macrophage** | 6 | 427 | 1 | 7 | 10 | 8 | 0 | 3 |
| **erythroid cell** | 2 | 16 | 176 | 483 | 250 | 482 | 470 | 766 |
| **erythroblast** | 3 | 98 | 691 | 1081 | 606 | 881 | 858 | 1427 |
| **erythrocyte** | 13 | 506 | 1488 | 1241 | 767 | 1196 | 1277 | 1768 |
| **basophil progenitor** | 0 | 0 | 2 | 4 | 2 | 6 | 5 | 6 |
| **basophil** | 0 | 1 | 34 | 60 | 35 | 61 | 51 | 64 |
| **fibroblast** | 0 | 4 | 3 | 4 | 9 | 22 | 4 | 18 |
| **dural fibroblast** | 16 | 917 | 1 | 14 | 5 | 3 | 2 | 2 |
| **endothelial cell** | 16 | 15 | 1 | 0 | 2 | 0 | 0 | 1 |
| **adipose-derived stromal cell** | 0 | 12 | 0 | 0 | 1 | 0 | 0 | 0 |
| **meningeal-Choroid Plexus cell** | 24 | 721 | 3 | 2 | 21 | 0 | 0 | 2 |
| **Omp+ cell** | 0 | 53 | 0 | 0 | 0 | 0 | 0 | 0 |
| **Gnb3+ cell** | 28 | 119 | 9 | 5 | 22 | 4 | 3 | 7 |
| **neuron** | 239 | 0 | 0 | 0 | 0 | 0 | 0 | 0 |
| **astrocyte** | 741 | 8 | 0 | 0 | 0 | 0 | 0 | 0 |
| **oligodendrocyte** | 110 | 1 | 0 | 0 | 0 | 0 | 0 | 0 |
| **megakaryocyte** | 1 | 0 | 1 | 1 | 2 | 4 | 0 | 3 |
| **innate lymphoid cell** | 1 | 37 | 5 | 6 | 2 | 6 | 3 | 2 |
