## Supplementary material for "Multi-omics and 3D-imaging reveal bone heterogeneity and unique calvaria cells in neuroinflammation": Table S2

significance pro and anti inflammatory scores

| | celltype | test | Il6 (pro) | Il1a (pro) | Il1b (pro) | Tnf (pro) | Ifng (pro) | Il11 (pro) | Il17d (pro) |
|---|---|---|---|---|---|---|---|---|---|
| 2 | B cell | Naive vs injury | 1 | 0.33442518 | 0.61158357 | 0.61158357 | 0.12980589 | 0.61158357 | 0.62005258 |
| 6 | B cell | MCAO Skull vs rest | 0.54043524 | 1 | 0.00047223 | 0.54043524 | 0.86832273 | 1 | 0.77068178 |
| 10 | B cell | Sham Skull vs rest | 1 | 1 | 0.95028513 | 1 | 1 | 1 | 1 |
| 14 | B cell | Naive Skull vs rest | 0.65652215 | 0.48613881 | 0.06071131 | 0.66200407 | 1 | 1 | 0.59947904 |
| 0 | all cells | Naive vs injury | 0.41620985 | 0.41620985 | 5.35E-14 | 1.70E-22 | 0.95895486 | 0.41620985 | 0.56651486 |
| 4 | all cells | MCAO Skull vs rest | 0.97694349 | 0.04210326 | 4.13E-121 | 5.47E-07 | 0.04028212 | 1 | 0.33417375 |
| 8 | all cells | Sham Skull vs rest | 0.00251069 | 1 | 4.64E-35 | 0.90284901 | 1 | 0.1482101 | 0.86925772 |
| 12 | all cells | Naive Skull vs rest | 0.30456216 | 0.56140624 | 8.62E-75 | 3.98E-12 | 0.30456216 | 0.01066663 | 0.66216144 |
| 3 | monocyte | Naive vs injury | 0.78179867 | 0.38787898 | 0.32030442 | 0.78179867 | 0.84497677 | 0.83706986 | 1 |
| 7 | monocyte | MCAO Skull vs rest | 0.90420911 | 0.54305431 | 9.03E-05 | 0.69226925 | 0.54305431 | 0.90420911 | 1 |
| 11 | monocyte | Sham Skull vs rest | 1 | 1 | 0.496398 | 1 | 1 | 1 | 1 |
| 15 | monocyte | Naive Skull vs rest | 0.39792297 | 1 | 0.00043619 | 1 | 1 | 1 | 1 |
| 1 | neutrophil | Naive vs injury | 0.24261038 | 0.51995706 | 1.46E-09 | 0.24261038 | 0.6919866 | 1 | 0.17779078 |
| 5 | neutrophil | MCAO Skull vs rest | 0.44953654 | 0.44953654 | 4.61E-124 | 7.58E-08 | 0.44953654 | 0.5136188 | 0.20318948 |
| 9 | neutrophil | Sham Skull vs rest | 0.41987436 | 0.77020652 | 8.75E-43 | 0.1145772 | 0.8230018 | 1 | 1 |
| 13 | neutrophil | Naive Skull vs rest | 0.59842008 | 0.56312767 | 4.94E-70 | 1.86E-10 | 0.49061778 | 1 | 1 |

| Il17f (pro) | Il18 (pro) | Il1rn (anti) | Tgfb1 (anti) | Il4 (anti) | Il10 (anti) | Il12a (anti) | Il13 (anti) | pro inflamma | anti inflammatory |
|---|---|---|---|---|---|---|---|---|---|
| 0.4882882 | 3.26E-06 | 0.04653797 | 1.63E-114 | 0.4882882 | 0.61158357 | 2.47E-76 | 1 | 6.40E-05 | 8.58E-126 |
| 0.90920751 | 1 | 1 | 0.7365592 | 1 | 0.97890787 | 0.7365592 | 1 | 0.86832273 | 0.7365592 |
| 1 | 1 | 1 | 0.08009127 | 1 | 1 | 1 | 1 | 1 | 1 |
| 0.59947904 | 0.80276337 | 0.59947904 | 0.0143556 | 1 | 0.48613881 | 0.16368318 | 1 | 1 | 0.03095693 |
| 0.95895486 | 1.27E-16 | 5.22E-181 | 0.0031787 | 0.76314061 | 0.76314061 | 4.47E-22 | 0.76314061 | 0.74262959 | 4.72E-50 |
| 0.97694349 | 0.97694349 | 0.97694349 | 6.65E-30 | 0.97694349 | 1 | 1 | 1 | 4.20E-47 | 1.14E-08 |
| 0.40056 | 3.50E-06 | 0.18868641 | 5.53E-16 | 4.15E-07 | 0.40056 | 1 | 1 | 7.48E-05 | 4.26E-07 |
| 0.30456216 | 0.04281961 | 6.27E-13 | 4.52E-08 | 0.67995584 | 0.37684958 | 0.04163971 | 0.66742226 | 3.45E-25 | 0.03035441 |
| 0.11475808 | 0.10981274 | 0.78179867 | 3.05E-45 | 1 | 0.38787898 | 1 | 1 | 0.78179867 | 5.98E-30 |
| 0.90420911 | 0.69226925 | 1 | 1 | 1 | 0.54305431 | 0.90420911 | 1 | 0.02812823 | 0.90420911 |
| 1 | 1 | 1 | 1 | 1 | 1 | 1 | 1 | 1 | 1 |
| 1 | 1 | 1 | 1 | 1 | 1 | 1 | 1 | 0.07568896 | 1 |
| 0.24261038 | 4.30E-10 | 0 | 1.54E-14 | 0.33005918 | 0.68246173 | 1.84E-09 | 0.57648865 | 2.09E-10 | 5.04E-243 |
| 0.01767457 | 0.30149221 | 0.7167755 | 5.48E-10 | 0.44953654 | 0.58367066 | 0.5136188 | 1 | 2.92E-106 | 0.10321416 |
| 1 | 0.25779343 | 0.00267161 | 1.26E-07 | 1 | 0.16205579 | 0.1145772 | 1 | 1.13E-27 | 0.41987436 |
| 0.49061778 | 0.49061778 | 1.09E-06 | 0.01949005 | 1 | 0.49061778 | 0.26063101 | 1 | 4.75E-63 | 0.01190291 |
