## Supplementary material for "Multi-omics and 3D-imaging reveal bone heterogeneity and unique calvaria cells in neuroinflammation": Table S3

Skull only genes MCAo condition

| | Gene names |
|---|---|
| 0 | Epha4 |
| 1 | Syn2 |
| 2 | Fap |
| 3 | Omd |
| 4 | Mmp2 |
| 5 | Mgp |
| 6 | Bhmt |
| 7 | Sptbn2 |
| 8 | Snap91 |
| 9 | Tnc |
| 10 | Stxbp1 |
| 11 | Fstl1 |
| 12 | Syn1 |
| 13 | Megf10 |
| 14 | B4gat1 |
| 15 | Ache |
