## Supplementary material for "Multi-omics and 3D-imaging reveal bone heterogeneity and unique calvaria cells in neuroinflammation": Table S4

| CODE | Age | Sex | SNP | SNP_CODE | Diagnose | GROUP _CODE | Motor_Area | Frontal_Parietal | Temporopolar | Skull_Base |
|---|---|---|---|---|---|---|---|---|---|---|
| PPMS-1 | 67.88 | 1 | MAB | 2 | PPMS | 1 | 0.903781372 | 0.652474864 | 1.287609757 | 1.342892541 |
| PPMS-2 | 58.88 | 1 | MAB | 2 | PPMS | 1 | 0.638061348 | 0.422751425 | 1.082257242 | 1.314877678 |
| PPMS-3 | 57.65 | 2 | HAB | 1 | PPMS | 1 | 1.15067046 | 1.270106712 | 0.831223654 | 0.874721652 |
| PPMS-4 | 58.21 | 2 | MAB | 2 | PPMS | 1 | 0.980155711 | 1.185095766 | 1.117128679 | 1.051389593 |
| PPMS-5 | 36.53 | 1 | unk. | 1.458 | PPMS | 1 | 1.720351675 | 1.446310278 | 1.387364217 | 1.637089351 |
| PPMS-6 | 35.44 | 1 | MAB | 2 | PPMS | 1 | 1.096225885 | 1.081854817 | 1.180655826 | 1.344410473 |
| PPMS-7 | 52.39 | 1 | HAB | 1 | PPMS | 1 | 1.381659918 | 1.912379968 | 1.018388111 | 1.145883732 |
| PPMS-8 | 40.87 | 2 | MAB | 2 | PPMS | 1 | 1.187506534 | 0.969528986 | 1.311495992 | 1.346946532 |
| PPMS-9 | 49.47 | 1 | HAB | 1 | PPMS | 1 | 1.077496012 | 1.22630181 | 1.07201261 | 1.271633873 |
| PPMS-10 | 54.62 | 2 | HAB | 1 | PPMS | 1 | 0.767448427 | 0.464762652 | 1.250458685 | 1.197819702 |
| PPMS-11 | 58.19 | 1 | MAB | 2 | PPMS | 1 | 0.893905323 | 0.758176855 | 1.101365777 | 1.273517413 |
| PPMS-12 | 48.35 | 1 | MAB | 2 | PPMS | 1 | 0.952308446 | 0.80795005 | 1.32111463 | 1.204569261 |
| PPMS-13 | 56.10 | 2 | unk. | 1.458 | PPMS | 1 | 0.880639178 | 0.492437912 | 1.337499624 | 1.464160539 |
| PPMS-14 | 50.45 | 1 | MAB | 2 | PPMS | 1 | 0.94977867 | 0.713570099 | 1.454161686 | 1.578809096 |
| RRMS-1 | 44.39 | 1 | HAB | 1 | RRMS | 2 | 0.709300767 | 0.471065876 | 1.315269304 | 1.171087604 |
| RRMS-2 | 44.47 | 2 | HAB | 1 | RRMS | 2 | 0.824336518 | 0.595506883 | 1.333078968 | 1.127031055 |
| RRMS-3 | 55.17 | 1 | HAB | 1 | RRMS | 2 | 0.960284592 | 0.852603427 | 1.221399903 | 1.365381368 |
| RRMS-4 | 33.57 | 2 | HAB | 1 | RRMS | 2 | 0.954612758 | 1.014248878 | 0.975331484 | 0.988815394 |
| RRMS-5 | 31.45 | 2 | HAB | 1 | RRMS | 2 | 0.776049726 | 0.770145915 | 1.038176762 | 1.01385608 |
| RRMS-6 | 52.75 | 1 | HAB | 1 | RRMS | 2 | 1.360959431 | 1.318053504 | 1.248993295 | 1.464950197 |
| RRMS-7 | 39.86 | 1 | HAB | 1 | PPMS | 2 | 0.921389184 | 0.897920148 | 0.976361524 | 1.054972457 |
| RRMS-8 | 39.57 | 2 | HAB | 1 | RRMS | 2 | 0.845734126 | 0.601918807 | 1.110205159 | 1.064128867 |
| RRMS-9 | 55.98 | 1 | MAB | 2 | RRMS | 2 | 0.929569575 | 0.694074733 | 1.45656532 | 1.266578944 |
| RRMS-10 | 26.78 | 1 | HAB | 1 | RRMS | 2 | 1.303853989 | 1.302381839 | 1.219134904 | 1.264856227 |
| RRMS-11 | 48.42 | 2 | HAB | 1 | RRMS | 2 | 1.073299389 | 1.192823975 | 0.999668826 | 1.230077885 |
| RRMS-12 | 50.26 | 2 | HAB | 1 | RRMS | 2 | 1.253624973 | 1.33026935 | 1.179584133 | 0.99670152 |
| RRMS-13 | 39.73 | 1 | MAB | 2 | RRMS | 2 | 1.791851093 | 1.580857302 | 1.298370422 | 1.568141898 |
| RRMS-14 | 47.58 | 2 | MAB | 2 | RRMS | 2 | 1.468411203 | 1.792361523 | 1.228309015 | 1.283930381 |
| RRMS-15 | 34.35 | 1 | MAB | 2 | RRMS | 2 | 0.963234599 | 0.85400887 | 0.984017657 | 1.293863602 |

| | | | | | | | | | | |
|---|---|---|---|---|---|---|---|---|---|---|
| Stroke-1 | 66.96 | 1 | HAB | 1 | Stroke | 3 | 0.854757069 | 0.779577506 | 1.059441194 | 0.87238866 |
| Stroke-2 | 57.88 | 1 | HAB | 1 | Stroke | 3 | 1.300828687 | 1.269585766 | 1.01897446 | 1.030897842 |
| Stroke-3 | 53.79 | 1 | HAB | 1 | Stroke | 3 | 1.488999683 | 1.489219257 | 1.315425166 | 1.128983024 |
| Stroke-4 | 63.10 | 1 | MAB | 2 | Stroke | 3 | 1.064943531 | 0.926583502 | 1.19694503 | 1.015361852 |
| Stroke-5 | 75.80 | 1 | HAB | 1 | Stroke | 3 | 0.944232201 | 0.989453376 | 1.12300469 | 1.119708093 |
| Stroke-6 | 69.70 | 1 | HAB | 1 | Stroke | 3 | 1.231330412 | 1.285846157 | 0.926133469 | 0.928002822 |
| Stroke-7 | 86.21 | 2 | HAB | 1 | Stroke | 3 | 0.787397416 | 0.747451779 | 1.290540243 | 0.858435926 |
| Stroke-8 | 82.84 | 1 | MAB | 2 | Stroke | 3 | 1.132765336 | 1.193974561 | 1.278277199 | 1.197838914 |
| Stroke-9 | 66.02 | 2 | MAB | 2 | Stroke | 3 | 2.121490271 | 1.712239695 | 1.757044273 | 1.157413628 |
| Stroke-10 | 75.07 | 2 | unk. | 1.458 | Stroke | 3 | 1.714356565 | 2.531680068 | 1.29927472 | 1.014127111 |

| | | | | | | | | | | |
|---|---|---|---|---|---|---|---|---|---|---|
| Control-1 | 55.77 | 2 | MAB | 2 | Control | 6 | 0.93613029 | 0.67589629 | 0.944912814 | 0.913689378 |
| Control-2 | 57.25 | 2 | MAB | 2 | Control | 6 | 1.383251653 | 1.984465029 | 0.821029476 | 0.79348867 |
| Control-3 | 67.26 | 1 | HAB | 1 | Control | 6 | 0.788870392 | 0.651801742 | 1.148107875 | 1.094179116 |
| Control-4 | 67.59 | 1 | MAB | 2 | Control | 6 | 0.774583679 | 0.631205292 | 0.868157929 | 0.90619328 |
| Control-5 | 69.10 | 2 | MAB | 2 | Control | 6 | 1.26651655 | 1.501710517 | 1.073519338 | 0.910749654 |
| Control-6 | 69.74 | 1 | MAB | 2 | Control | 6 | 0.86695111 | 0.569000634 | 0.99354911 | 1.091112797 |
| Control-7 | 71.26 | 1 | MAB | 2 | Control | 6 | 0.87715914 | 0.770552224 | 1.163284452 | 1.13463507 |
| Control-8 | 71.41 | 2 | MAB | 2 | Control | 6 | 0.965859095 | 0.795855146 | 0.939397646 | 0.898335036 |
| Control-9 | 72.59 | 2 | MAB | 2 | Control | 6 | 0.979919601 | 0.87006836 | 1.033743735 | 1.012741129 |
| Control-10 | 72.21 | 1 | HAB | 1 | Control | 6 | 0.70900339 | 0.576433976 | 0.98541207 | 0.994563977 |
| Control-11 | 72.08 | 2 | MAB | 2 | Control | 6 | 1.230800491 | 1.207871122 | 1.078774131 | 1.266566143 |
| Control-12 | 72.13 | 1 | HAB | 1 | Control | 6 | 1.30744972 | 1.936258562 | 0.918935731 | 0.831901784 |
| Control-13 | 74.85 | 2 | MAB | 2 | Control | 6 | 1.135542284 | 1.187278833 | 0.878653918 | 0.96709695 |
| Control-14 | 77.04 | 1 | MAB | 2 | Control | 6 | 0.999500684 | 0.995611632 | 1.183935136 | 1.203732249 |
| Control-15 | 78.39 | 1 | unk. | 1.458 | Control | 6 | 0.872322042 | 0.948616109 | 0.962672 | 0.980355339 |
| Control-16 | 79.89 | 1 | HAB | 1 | Control | 6 | 0.967505726 | 0.973371833 | 0.946383813 | 1.007232701 |
| Control-17 | 80.71 | 2 | MAB | 2 | Control | 6 | 0.938634151 | 0.724002698 | 1.059530826 | 0.993426729 |

| | | | | | | | | | | |
|---|---|---|---|---|---|---|---|---|---|---|
| Control-18 | 24.27 | 2 | MAB | 2 | Control | 6 | 0.621747574 | 0.808667295 | 0.632909513 | 0.847426524 |
| Control-19 | 59.92 | 2 | MAB | 2 | Control | 6 | 1.081777372 | 1.057706838 | 1.139189988 | 1.076233732 |
| Control-20 | 24.84 | 2 | HAB | 1 | Control | 6 | 0.789032634 | 1.024235658 | 1.120617028 | 1.079227083 |
| Control-21 | 23.97 | 2 | MAB | 2 | Control | 6 | 1.327681884 | 1.382687953 | 0.996543002 | 1.037681663 |
| Control-22 | 26.08 | 2 | HAB | 1 | Control | 6 | 1.542453101 | 1.620861018 | 1.119114581 | 1.025192009 |

| | | | | | | | | | | |
|---|---|---|---|---|---|---|---|---|---|---|
| Control-23 | 21.38 | 2 | HAB | 1 | Control | 6 | 1.501759516 | 1.672322679 | 0.979098607 | 1.003139779 |
| Control-24 | 26.82 | 1 | MAB | 2 | Control | 6 | 0.951111584 | 0.757014803 | 1.069854415 | 1.181466601 |
| Control-25 | 54.22 | 2 | MAB | 2 | Control | 6 | 0.996398772 | 0.76203104 | 1.089177612 | 1.083744603 |
| Control-26 | 39.54 | 2 | unk. | 1.458 | Control | 6 | 0.623660548 | 0.543070156 | 0.846925122 | 0.834084463 |
| Control-27 | 69.52 | 1 | HAB | 1 | Control | 6 | 0.564377013 | 0.37140256 | 1.006570132 | 0.831803543 |
| 4RT-1 | 57.64 | 1 | HAB | 1 | 4RT | 4 | 1.15449454 | 1.091742907 | 1.165020866 | 1.161795011 |
| 4RT-2 | 62.98 | 1 | HAB | 1 | 4RT | 4 | 1.048100342 | 0.979492768 | 1.066640973 | 0.996847001 |
| 4RT-3 | 53.67 | 2 | MAB | 2 | 4RT | 4 | 1.046847161 | 1.425070948 | 0.969919967 | 0.88755932 |
| 4RT-4 | 66.11 | 2 | HAB | 1 | 4RT | 4 | 1.070056698 | 1.078585147 | 0.840634773 | 0.760173099 |
| 4RT-5 | 78.61 | 2 | MAB | 2 | 4RT | 4 | 0.900843315 | 0.849960648 | 0.910154269 | 0.842209203 |
| 4RT-6 | 60.96 | 2 | HAB | 1 | 4RT | 4 | 1.40057355 | 1.620229017 | 1.100973015 | 1.099492662 |
| 4RT-7 | 72.05 | 1 | HAB | 1 | 4RT | 4 | 1.136009245 | 0.890903284 | 1.181035193 | 1.180606079 |
| 4RT-8 | 72.83 | 1 | HAB | 1 | 4RT | 4 | 1.085488419 | 0.92345775 | 1.238056548 | 1.148859734 |
| 4RT-9 | 65.74 | 2 | MAB | 2 | 4RT | 4 | 1.821575501 | 1.965431009 | 1.022848014 | 1.187131503 |
| 4RT-10 | 52.00 | 2 | MAB | 2 | 4RT | 4 | 1.119207791 | 1.135607304 | 1.084820519 | 1.244185091 |
| 4RT-11 | 64.61 | 2 | HAB | 1 | 4RT | 4 | 1.433034265 | 1.870498256 | 1.225509803 | 1.150574339 |
| 4RT-12 | 82.91 | 1 | HAB | 1 | 4RT | 4 | 0.934836546 | 0.849444638 | 1.262814785 | 1.118243499 |
| 4RT-13 | 70.05 | 1 | MAB | 2 | 4RT | 4 | 1.089690154 | 1.307913675 | 1.155135229 | 1.202347498 |
| 4RT-14 | 82.20 | 2 | HAB | 1 | 4RT | 4 | 1.337987774 | 1.10140985 | 1.042970252 | 0.996013679 |
| 4RT-15 | 77.57 | 2 | MAB | 2 | 4RT | 4 | 1.972535604 | 2.36950774 | 1.1413868 | 1.094842393 |
| 4RT-16 | 63.53 | 1 | HAB | 1 | 4RT | 4 | 1.097223518 | 0.961611729 | 1.083592343 | 1.083376295 |
| 4RT-17 | 78.72 | 1 | MAB | 2 | 4RT | 4 | 1.016578559 | 0.835509531 | 0.938907672 | 0.966578143 |
| 4RT-18 | 58.99 | 1 | HAB | 1 | 4RT | 4 | 1.278626313 | 1.180783516 | 0.898435536 | 0.895970222 |
| 4RT-19 | 69.96 | 2 | HAB | 1 | 4RT | 4 | 0.888530466 | 0.879457061 | 1.029801539 | 1.037387716 |
| 4RT-20 | 68.98 | 1 | MAB | 2 | 4RT | 4 | 0.766249946 | 0.737408644 | 1.078374288 | 1.208756003 |
| 4RT-21 | 65.60 | 2 | HAB | 1 | 4RT | 4 | 1.769133777 | 1.739105825 | 1.054767173 | 0.999987451 |
| 4RT-22 | 73.43 | 2 | HAB | 1 | 4RT | 4 | 0.822658343 | 0.991967218 | 0.908260194 | 0.794624369 |
| 4RT-23 | 68.54 | 2 | HAB | 1 | 4RT | 4 | 0.939712737 | 0.972932828 | 1.008823384 | 0.847289819 |
| 4RT-24 | 75.46 | 2 | HAB | 1 | 4RT | 4 | 0.983120133 | 1.030040737 | 0.919660134 | 1.055967126 |
| 4RT-25 | 72.74 | 1 | HAB | 1 | 4RT | 4 | 1.367014056 | 1.306634526 | 0.96743466 | 1.032897825 |
| 4RT-26 | 74.25 | 2 | HAB | 1 | 4RT | 4 | 1.194727302 | 1.071824333 | 0.989241501 | 1.00230545 |
| 4RT-27 | 67.25 | 1 | HAB | 1 | 4RT | 4 | 0.930578947 | 1.085369644 | 0.9340843 | 0.89042301 |
| 4RT-28 | 75.83 | 1 | MAB | 2 | 4RT | 4 | 1.333028798 | 1.121718447 | 1.191096793 | 1.111884947 |

| | | | | | | | | | |
|---|---|---|---|---|---|---|---|---|---|
| 4RT-29 | 83.83 | 1 | HAB | 1 | 4RT | 4 | 1.168050919 | 1.439978421 | 1.191084531 | 1.167602807 |
| 4RT-30 | 62.91 | 1 | MAB | 2 | 4RT | 4 | 1.450055868 | 1.255215179 | 1.08865511 | 1.298428614 |
| 4RT-31 | 66.28 | 2 | MAB | 2 | 4RT | 4 | 1.299920953 | 1.700167072 | 1.137640785 | 1.015917002 |
| 4RT-32 | 70.10 | 2 | HAB | 1 | 4RT | 4 | 0.88792314 | 0.96374162 | 0.859331644 | 0.720488794 |
| 4RT-33 | 74.63 | 1 | HAB | 1 | 4RT | 4 | 1.294685316 | 1.520403565 | 0.824274485 | 0.793193906 |
| 4RT-34 | 75.06 | 2 | HAB | 1 | 4RT | 4 | 0.937531772 | 1.057196743 | 1.04164541 | 1.059322658 |
| 4RT-35 | 56.81 | 1 | HAB | 1 | 4RT | 4 | 1.262185867 | 1.076898695 | 1.086322995 | 1.175765633 |
| 4RT-36 | 64.52 | 1 | HAB | 1 | 4RT | 4 | 2.023467213 | 2.642136854 | 1.191795169 | 1.055716727 |
| 4RT-37 | 86.23 | 1 | MAB | 2 | 4RT | 4 | 0.840997426 | 0.805567192 | 0.891736239 | 1.021813238 |
| 4RT-38 | 56.07 | 2 | HAB | 1 | 4RT | 4 | 1.262922942 | 1.213032238 | 1.087716729 | 0.890170805 |
| 4RT-39 | 74.87 | 2 | HAB | 1 | 4RT | 4 | 0.866406707 | 0.890606372 | 1.016992745 | 0.880520182 |
| 4RT-40 | 66.73 | 1 | HAB | 1 | 4RT | 4 | 0.876071254 | 0.898238701 | 0.877304995 | 0.813955468 |
| 4RT-41 | 76.96 | 1 | HAB | 1 | 4RT | 4 | 0.976080867 | 0.936987679 | 1.260620483 | 1.171758983 |
| 4RT-42 | 66.70 | 2 | HAB | 1 | 4RT | 4 | 1.252611913 | 1.489263713 | 0.839794735 | 0.891207472 |
| 4RT-43 | 68.90 | 2 | HAB | 1 | 4RT | 4 | 1.495798328 | 1.170313605 | 0.91751043 | 0.931705264 |
| AD-1 | 78.51 | 2 | MAB | 2 | AD | 5 | 1.156784491 | 1.395244118 | 1.099022404 | 0.98164681 |
| AD-2 | 67.77 | 2 | MAB | 2 | AD | 5 | 1.081138448 | 1.360504071 | 1.306875671 | 1.336979934 |
| AD-3 | 72.70 | 2 | MAB | 2 | AD | 5 | 0.796952952 | 0.954566459 | 0.970605831 | 0.861088653 |
| AD-4 | 74.52 | 2 | MAB | 2 | AD | 5 | 1.103414432 | 1.285507368 | 1.019319454 | 1.093608048 |
| AD-5 | 62.68 | 2 | HAB | 1 | AD | 5 | 1.351869228 | 1.117248106 | 1.008309082 | 0.96018398 |
| AD-6 | 64.73 | 2 | MAB | 2 | AD | 5 | 1.390689294 | 1.990557763 | 1.171646041 | 0.95366958 |
| AD-7 | 57.09 | 2 | HAB | 1 | AD | 5 | 1.641841124 | 1.516728501 | 0.975227934 | 0.783108696 |
| AD-8 | 80.00 | 2 | MAB | 2 | AD | 5 | 1.546824281 | 1.983981525 | 1.155752968 | 1.108559656 |
| AD-9 | 73.92 | 2 | HAB | 1 | AD | 5 | 1.042291731 | 1.057329056 | 1.115644355 | 1.108271276 |
| AD-10 | 59.22 | 2 | MAB | 2 | AD | 5 | 1.04902544 | 0.887850707 | 1.084276008 | 0.963525534 |
| AD-11 | 75.65 | 2 | HAB | 1 | AD | 5 | 1.234668855 | 1.249231674 | 0.929397973 | 0.893190194 |
| AD-12 | 74.92 | 2 | MAB | 2 | AD | 5 | 1.181921099 | 1.417748819 | 0.986114423 | 1.003039084 |
| AD-13 | 81.41 | 2 | MAB | 2 | AD | 5 | 1.262879625 | 1.682230912 | 0.930016672 | 0.834350469 |
| AD-14 | 69.14 | 2 | HAB | 1 | AD | 5 | 0.761435748 | 0.790345315 | 0.845372362 | 0.809198624 |
| AD-15 | 82.30 | 2 | HAB | 1 | AD | 5 | 1.31583847 | 1.39999858 | 1.079331181 | 1.07467845 |
| AD-16 | 53.33 | 2 | MAB | 2 | AD | 5 | 1.174472063 | 1.097295203 | 1.038019939 | 1.088596022 |
| AD-17 | 58.58 | 2 | MAB | 2 | AD | 5 | 1.753814598 | 2.571592208 | 0.739953376 | 0.970040378 |
| AD-18 | 66.12 | 2 | MAB | 2 | AD | 5 | 1.358908149 | 1.330686706 | 1.112773596 | 1.205550794 |

| | | | | | | | | | | |
|---|---|---|---|---|---|---|---|---|---|---|
| AD-19 | 65.87 | 2 | MAB | 2 | AD | 5 | 1.391480026 | 2.256282978 | 0.748134136 | 0.863119264 |
| AD-20 | 80.43 | 2 | MAB | 2 | AD | 5 | 1.327946472 | 1.686479815 | 0.932577995 | 0.982235794 |
| AD-21 | 63.13 | 2 | MAB | 2 | AD | 5 | 1.818740976 | 2.47943052 | 1.159368261 | 1.219119632 |
| AD-22 | 74.07 | 2 | HAB | 1 | AD | 5 | 1.456358105 | 2.101809789 | 0.963115753 | 0.950745849 |
| AD-23 | 74.16 | 2 | HAB | 1 | AD | 5 | 1.086851604 | 1.195830659 | 1.160394119 | 1.013571302 |
| AD-24 | 69.01 | 2 | HAB | 1 | AD | 5 | 0.941976868 | 1.12059851 | 0.899359976 | 0.79532134 |
| AD-25 | 74.64 | 2 | MAB | 2 | AD | 5 | 0.997201273 | 0.841102458 | 1.308862426 | 1.034432466 |
| AD-26 | 64.73 | 2 | MAB | 2 | AD | 5 | 1.639355645 | 2.38055107 | 1.156938647 | 1.009682694 |
| AD-27 | 53.82 | 2 | MAB | 2 | AD | 5 | 1.512411899 | 1.650549944 | 1.006009622 | 1.022590425 |
| AD-28 | 77.93 | 2 | HAB | 1 | AD | 5 | 1.315046766 | 1.741286439 | 0.806527216 | 0.859529191 |
| AD-29 | 67.24 | 2 | HAB | 1 | AD | 5 | 1.128331599 | 1.553536706 | 0.987779745 | 0.667134508 |
| AD-30 | 62.49 | 1 | MAB | 2 | AD | 5 | 1.392142041 | 1.326453128 | 1.018724995 | 1.067326017 |
| AD-31 | 66.56 | 1 | HAB | 1 | AD | 5 | 0.750827329 | 0.652187361 | 0.869175077 | 0.97768804 |
| AD-32 | 80.70 | 1 | HAB | 1 | AD | 5 | 0.734812979 | 0.604575819 | 0.946290141 | 0.973705348 |
| AD-33 | 72.07 | 1 | HAB | 1 | AD | 5 | 0.961736826 | 0.813427321 | 0.998743884 | 1.013543388 |
| AD-34 | 72.58 | 1 | HAB | 1 | AD | 5 | 0.875919264 | 1.049187303 | 1.100471417 | 0.986660281 |
| AD-35 | 76.98 | 1 | MAB | 2 | AD | 5 | 1.083848588 | 1.025131623 | 1.136592371 | 1.279296584 |
| AD-36 | 72.01 | 1 | HAB | 1 | AD | 5 | 0.818180682 | 0.839660101 | 0.811978729 | 0.914973175 |
| AD-37 | 78.08 | 1 | HAB | 1 | AD | 5 | 1.044237414 | 0.915635509 | 0.865375787 | 1.111715186 |
| AD-38 | 69.56 | 1 | unk. | 1.458 | AD | 5 | 1.072323256 | 0.850849746 | 1.139909261 | 1.064846611 |
| AD-39 | 67.39 | 1 | MAB | 2 | AD | 5 | 1.802509375 | 1.752039092 | 1.224140285 | 1.165310128 |
| AD-40 | 77.75 | 1 | MAB | 2 | AD | 5 | 0.870857225 | 1.005819283 | 0.94964899 | 0.967604714 |
| AD-41 | 75.72 | 1 | MAB | 2 | AD | 5 | 1.073014702 | 1.029844054 | 1.004739421 | 1.059950087 |
| AD-42 | 77.74 | 1 | MAB | 2 | AD | 5 | 1.094444041 | 1.008496324 | 1.089323594 | 1.143491557 |
| AD-43 | 79.44 | 1 | MAB | 2 | AD | 5 | 0.967321101 | 0.78733347 | 1.097525515 | 1.102245035 |
| AD-44 | 74.57 | 1 | HAB | 1 | AD | 5 | 0.714844426 | 0.606299382 | 0.775923047 | 0.873887232 |
| AD-45 | 70.48 | 1 | HAB | 1 | AD | 5 | 0.855268765 | 0.682762369 | 1.129673589 | 1.087061226 |
| AD-46 | 60.30 | 1 | HAB | 1 | AD | 5 | 1.403591794 | 1.536612188 | 1.025922984 | 1.075677148 |
| AD-47 | 81.35 | 1 | HAB | 1 | AD | 5 | 0.844703855 | 0.763089166 | 0.833912648 | 0.824454883 |
| AD-48 | 72.36 | 1 | HAB | 1 | AD | 5 | 1.111822755 | 0.955204418 | 0.82911993 | 0.994485821 |
| AD-49 | 79.65 | 1 | MAB | 2 | AD | 5 | 0.892288979 | 0.825984833 | 0.958870025 | 0.909856617 |
| AD-50 | 70.77 | 2 | HAB | 1 | AD | 5 | 1.699230136 | 1.551450199 | 0.948732904 | 1.003350839 |
